# Single amino acid exchange in ACTIN2 confers increased tolerance to oxidative stress in Arabidopsis *der1-3* mutant

**DOI:** 10.1101/2020.12.17.423225

**Authors:** Lenka Kuběnová, Tomáš Takáč, Jozef Šamaj, Miroslav Ovečka

## Abstract

Single-point mutation in the *ACTIN2* gene of *der1-3* mutant revealed that ACTIN2 is an essential actin isovariant required for root hair tip growth, and leads to shorter, thinner and more randomly oriented actin filaments in comparison to wild-type C24 genotype. Actin cytoskeleton has been linked to plant defence against oxidative stress, but it is not clear how altered structural organization and dynamics of actin filaments may help plants to cope with oxidative stress. In this study, we characterized seed germination, root growth, plant biomass, actin organization and antioxidant activity of *der1-3* mutant under oxidative stress induced by paraquat and H_2_O_2_. Under these conditions, plant growth was better in *der1-3* mutant, while actin cytoskeleton in *der1-3* carrying *pro35S::GFP:FABD2* construct showed lower bundling rate and higher dynamicity. Biochemical analyses documented lower degree of lipid peroxidation, elevated capacity to decompose superoxide and hydrogen peroxide. These results support the view that *der1-3* mutant is more resistant to oxidative stress. Single amino acid exchange in mutated ACTIN2 protein (Cys to Arg at the position 97) is topologically exposed to the protein surface and we propose that this might alter protein post-translational modifications and/or protein-protein interactions, leading to enhanced tolerance of *der1-3* mutant against oxidative stress.

**Highlight:** Topological position of one amino acid exchanged in the ACTIN2 protein structure in *der1-3* mutant enhanced tolerance to oxidative stress through increased capacity to decompose reactive oxygen species, lower bundling and enhanced dynamicity of the actin cytoskeleton.

## Introduction

Plants are continuously exposed to fluctuating environmental conditions, including adverse biotic and abiotic stressors. Oxidative stress, alone or in combination with other stress factors, may significantly disrupt normal cellular homeostasis in plants. Oxidative stress significantly increases symplastic and apoplastic amount of reactive oxygen species (ROS) such as superoxide (O_2_^•−^), hydrogen peroxide (H_2_O_2_), hydroxyl radical (OH^•^) and singlet oxygen (^1^O_2_). Although ROS serve also as signalling molecules, playing important roles in the regulation of numerous plant developmental processes (Mittler *et al*., 2011; Baxter *et al*., 2014; Mhamdi and Van Breusegem, 2018), they are generated as toxic by-products of the aerobic metabolisms under stress conditions (Konig *et al*., 2012; Foyer and Noctor, 2013; Vaahtera *et al*., 2014; Mignolet-Spruyt *et al*., 2016; Mittler *et al*., 2017). In general, production of ROS with metabolic or stress-related origin, is controlled by components of redox signalling pathways. These maintain cellular ROS homeostasis, since both low and high ROS levels are undesirable for plant cells. Thus, an equilibrated threshold of ROS is maintained and controlled by the activity of antioxidant enzymes from the family of superoxide dismutases (SODs), catalases, peroxidases, gluthatione peroxidases, iron uptake/storage regulating mechanisms, and a network of thio- and glutaredoxins (Vanderauwera *et al*., 2011; Mittler, 2017).

Common approach inducing oxidative stress in plants experimentally is based on external application of paraquat (PQ, 1, 1’-dimethyl-4, 4’-bipyridinium chloride) and hydrogen peroxide (H_2_O_2_). PQ, or methyl viologen, is widely used herbicide, which passes rapidly to the cells (Riley *et al*., 1976; Hawkes, 2014). PQ is effective particularly in photosynthetically-active plant tissues (Krieger-Liszkay *et al*., 2011). Primary place of its activity is chloroplast, where PQ takes away electrons, probably from photosystem I and ferredoxin. This process leads to the formation of stable reduced cationic radical, reacting rapidly with molecular oxygen to form superoxides (Farrington *et al*., 1973; Bus *et al*., 1974). Superoxide anion interferes with antioxidant defence mechanisms leading to the damage of cells due to numerous chain reactions (Kunert and Dodge, 1989). Treatment of plants with PQ affects also gene expression. Alterations have been documented in the expression of numerous genes encoding different protein kinases (RLKs, MAPKs and CDPKs), antioxidant enzymes like ascorbate peroxidase, CuZn SOD (CSD), FeSOD (FSD), some transcription factors (MYB, MYC) or in genes securing cell structural integrity (Han *et al*., 2014). All these changes likewise lead to significant oxidation of proteins or nucleic acids (Xiong *et al.,* 2007). Typical phenotypical reaction of affected plants is wilting and chlorosis. Prolonged PQ exposure causes browning of damaged tissues and severe chlorosis, leading to falling off the leaves (Hawkes, 2014).

H_2_O_2_ is generated in chloroplasts, mitochondria and peroxisomes as inevitable by-product of aerobic metabolism, or it is produced under stress conditions (both biotic and abiotic stress) (Maurino and Flügge, 2008). It is a relatively long-living molecule (up to 1 ms) with the ability to pass membranes either by diffusion of actively via aquaporins (Levine *et al*., 1994; Willekens *et al*., 1997; Dat *et al*., 2000; Miller *et al*., 2010). The biological activity of H_2_O_2_ is mediated by its ability to oxidize free SH groups (Dat *et al*., 2000). Excessive exogenous H_2_O_2_ application induces rapid cell death and necrosis of plant cells, even without passing through the apoptotic stage (O’Brien *et al*., 1998; Yao *at al*., 2001; Maurino and Flügge, 2008). External application of H_2_O_2_ can stimulate some morphogenetic events in plants, like adventitious root initiation in flax hypocotyls (Takáč *et al*., 2016). H_2_O_2_, particularly at higher concentrations, affects the expression pattern of genes involved in diverse plant defence responses. It can be exemplified by changing expression patterns of genes encoding biosynthetic enzymes of phenylpropanoids, lignin and salicylic acid (Douglas 1996; Mauch-Mani and Slusarenko, 1996), as well as enzymes protective against oxidative stress, like glutathione S-transferase (GST; Pickett and Lu, 1989) and anthranilate synthase (ASA1), which is required for the biosynthesis of the phytoalexin camalexin (Desikan *et al*., 1998). Excessive formation of H_2_O_2_ causes also oxidative impairments of photosynthetic apparatus (Park *et al*., 1998; Rao and Davis, 1999; Dat *et al*., 2000).

Actin filaments are essential cytoskeletal components, playing important roles in cellular (e.g. cytoplasmic streaming, organelle movement, vesicular trafficking) and developmental processes (e.g. establishment and maintenance of cell polarity and shape, cell division plane determination, tip growth). Typical feature of actin cytoskeleton is its ability to perform dynamic structural reorganizations (Wasteneys and Galway, 2003; Staiger and Blanchoin, 2006). The actin cytoskeleton is also involved in signalling events triggered by diverse external stimuli. Among others, actin cytoskeleton remodelling is part of abiotic stress response mechanisms in plants (Zhou *et al*., 2010). Interestingly, dysfunctional ACTIN2 isoform or destabilization of actin microfilaments using cytochalasin D alter localization of *A. thaliana* Respiratory burst oxidase homolog protein C (AtRbohC) during root hair development in Arabidopsis (Takeda *et al*., 2008). Connection between intracellular distribution pattern of NDC1 (NAD(P)H dehydrogenases type II), able to reduce mitochondrial ROS production and ACTIN2 was also revealed (Wallström *et al*., 2012). These data suggest a supporting role of actin filaments in the mediation of both short- and long-term plant responses to oxidative stress conditions. The dependence between changing dynamics of the actin cytoskeleton and elevated ROS level was described in Arabidopsis root tip cells under salt stress. Treatment with NADPH oxidase inhibitor diphenyleneiodonium prevented salt stress-induced ROS increase, but treatment with actin inhibitors latrunculin B or jasplakinolide caused enhanced ROS accumulation in salt stress-treated root cells (Liu *et al*., 2012). Actin microfilaments play an important role in vesicular trafficking, which is linking ROS signalling with auxin transport (Zwiewka *et al*., 2019). In proposed model, oxidative stress caused by H_2_O_2_ affects dynamics of actin cytoskeleton, which subsequently interferes with ARF-GEF-dependent trafficking of PIN2 from the plasma membrane to early endosomes (Zwiewka *et al*., 2019). However, the complete mechanism linking structural and dynamic properties of actin cytoskeleton to oxidative stress in plants is not fully understood yet.

The actin cytoskeleton is essential for tip growth of root hairs. Indispensable functions of actin in growing root hairs were documented by pharmacological (Baluška *et al*., 2000) and genetic (Gilliland *et al*., 2002; Ringli *et al*., 2002) means. Land-plant evolution brought a diversification into reproductive and vegetative classes of actin, later ones represented by ACT2, ACT7 and ACT8 (McDowell *et al*., 1996; Meagher *et al*., 1999). Genetic approaches using chemically-induced single-point mutation (Ringli *et al*., 2002) or insertional knockout mutation (Gilliland *et al*., 2002) revealed that *ACT2* is essential for proper root hair tip growth. Taking into account that the expression level of *ACT2* gene is not affected by the single point mutations in the *DER1* locus and wild-type levels of *ACT2* expression has been documented in the *der1* (*deformed root hairs1*) mutants (Ringli *et al*., 2002), the palette of received mutants (*der1-1*, *der1-2*, *der1-3*) showed different degrees of the mutant root hair phenotype (Ringli *et al*., 2002). It provides an opportunity to characterize involvement of single-point mutation in partially functional ACT2 protein in different aspects of plant development (Vaškebová *et al*., 2018). In this study, we describe growth and developmental parameters of *der1-3* mutant, bearing strong *ACT2* mutation phenotype (Ringli *et al*., 2002), under oxidative stress caused by PQ and H_2_O_2_. In comparison to plants of wild-type (C24 genotype), post-germination root growth, biomass production, antioxidant activity and prevention of lipid peroxidation were more effective in *der1-3* mutant. Considering lower bundling rate and higher dynamicity of the actin cytoskeleton, we conclude that *der1-3* mutant plants are more resistant to mild and severe oxidative stress induced by PQ or H_2_O_2_ in the culture medium.

## Materials and methods

### Plant material and cultivation in vitro

Seeds of *Arabidopsis thaliana* (L.) Heynh., ecotype C24, *der1-3* mutant and transgenic lines expressing marker for visualization of the actin cytoskeleton were surface sterilized and planted into ½ Murasighe and Skoog medium without vitamins solidified with 0.6 % Gellan gum (Alfa Aesar, ThermoFisher). Seeds on medium in Petri dishes were stratified at 4°C for 3 days for synchronized germination. After stratification, seeds on plates were cultivated *in vitro* vertically in culture chamber at 21°C, 70 % humidity, and 16/8h light/dark cycle.

### Transgenic lines and transformation method

Plants of *Arabidopsis thaliana* (L.) Heynh. ecotype C24 (Beemster *et al*., 2002) and *der1-3* mutant (Ringli *et al*., 2002) were transformed with *Agrobacterium tumefaciens* stain GV3101 carrying a construct *pro35S::GFP:FABD2*, coding for F-actin binding domain 2 of Arabidopsis FIMBRIN 1 (FABD2) fused to green fluorescent protein (GFP; Voigt *et al*., 2005). These lines were used for fluorescent visualization of actin filaments (Vaškebová *et al*., 2018). Briefly, this construct was prepared in pCB302 vector with rifampicin and kanamycin resistance by classical cloning method with herbicide phosphinothricin as the selection marker *in planta*. Stable transformation was used according to Clough and Bent (1998). Plants (3-4 weeks old) were soaked in *Agrobacterium tumefaciens* cultures for 10 seconds and were stabilized in dark overnight. After that, plants were cultivated in culture chamber at 24°C, 60 % humidity, 16/8h light/dark photoperiod. Transformation was repeated after one week. Seeds of T_1_ generation were planted for selection on ½ MS media with phosphinothricin (50 mg.ml^-1^). Transgenic plants were selected for the presence of GFP fusion proteins using epifluorescence zoom microscope Axio Zoom.V16 (Carl Zeiss, Germany). For further experiments, seeds of T_3_ generation were used.

### Application of stress factors

Oxidative stress was induced by adding of three different concentrations of paraquat (PQ; 0.1; 0.2 and 0.5 μmol.l^-1^), and four different concentrations of H_2_O_2_ (0.5; 1; 1.5 and 3 mmol.l^-1^) to culture medium. Seeds were planted either directly on ½ MS media containing different concentrations of PQ, or 3 days old plants germinated on control media were transferred to media containing different concentrations of PQ or H_2_O_2_.

### Phenotypical analysis

Plants germinating on control media growing *in vitro* were scanned directly on plates every 24 hours for 11 days from the day of germination. Plants germinating on control media and transferred to stress conditions were scanned on plates every 24 hours for additional 4 days after their transfer. Images from the scanner (Image Scanner III, GE Healthcare, Chicago, IL, USA) were used for measurement of the primary root length. Images documenting phenotype of plants growing in plates were prepared with Nikon 7000 camera equipped with macro-objective Sigma 50 mm (2.8 focal distance) in time points indicated in the corresponding figure captions. Fresh weights of separated shoots and roots were measured from 18 days old plants growing on media containing PQ.

### Sample preparation and microscopic analysis

Samples for microscopic analysis were prepared in microscopic chambers filled with liquid culture medium according to Ovečka *et al*. (2005). Oxidative stress was induced using liquid culture medium containing 0.1 μmol.l^-1^ of PQ. Samples were firstly observed under the microscope in control medium for 30 min and then medium containing PQ was applied using perfusion of microscopic chamber. Total volume of the medium applied was 100 μl, added by perfusion sequentially 10 times with 10 μl. After perfusion, plants were carefully covered with parafilm and samples were scanned in the microscope every 30 s for further 30 min. Live cell imaging of actin cytoskeleton in hypocotyl epidermal cells of C24 ecotype and *der1-3* mutant expressing a construct *pro35S::GFP:FABD2* was performed in a fast scanning mode using spinning disk microscope Cell Observer SD Axio Observer Z1 (Carl Zeiss, Germany), equipped with EC Plan-Neofluar 40×/1.3 NA oil immersion objective (Carl Zeiss, Germany) and Plan-Apochromat 63×/1.4 NA oil immersion objective (Carl Zeiss, Germany). Samples were imaged with excitation laser line 488 nm and emission filter BP525/50. Laser power was set up not to exceed 50% of the laser intensity range available. Samples were scanned in a Z-stack mode in a time range of every 30 seconds for 30 min. Images were acquired with the Evolve 512 EM CCD camera with the exposure time 500-750 ms per optical section. Orthogonal projections of 6 to 10 optical sections from Z-stacks were used for preparation of videos and measurement of actin filament skewness and occupancy. Semiquantitative analysis of actin filament dynamics in hypocotyl epidermal cells was presented by pseudocolouring displacement analysis. Images were acquired at the beginning, after 15 min and after 30 min of time-point scanning, individually coloured red, green, and blue, respectively, and merged. Overlay of all three colours creating a white one indicated lowering, or eventually stopping, of the actin dynamic activity.

### Histochemical detection of O2^•−^ and H_2_O_2_ production

Plants (3 days old) were transferred from control media to media containing 0.1 μmol.l^-1^ PQ and 3 mmol.l^-1^ H_2_O_2_ and histochemical detection of ROS was done 11 days after the transfer (plants were 14 days old). Superoxide (O2^•−^) was detected by NBT (nitrotetrazolium blue) staining according Ramel *et al*. (2009). H_2_O_2_ detection was done with DAB (diaminobenzidine) staining according Daudi *et al*. (2012). Plants after staining were mounted and imaged in Axio Zoom.V16 (Carl Zeiss, Germany). Staining intensity mean values in cotyledons and leaves were measured and quantified in ZEN 2 (blue edition; Carl Zeiss, Germany) software.

### Analysis of enzymatic activity and immunoblotting

For superoxide dismutase (SOD) activity examination, proteins were extracted using Na-phosphate extraction buffer containing 50 mM Na-phosphate buffer (pH 7.8), 2 mM EDTA, 2 mM ascorbic acid and 10% (v/v) glycerol. SOD activities were visualised on native PAGE gels as described by Takáč *et al*. (2014). For immunoblotting, the enzyme extracts were enriched with 4x Laemmli SDS buffer (to reach final concentration of 10% v/v glycerol, 60 mM Tris/HCl pH 6.8, 2% w/v SDS, 0.002% w/v bromophenol blue and 5% v/v β-mercaptoethanol). Afterwards, the samples were boiled at 95°C for 5 min. Equal amounts of proteins (15 μg) were loaded on 10% SDS PAGE gels. Immunoblotting analysis and chemiluminiscence signal development were carried out according to Takáč *et al*. (2017). As a primary antibodies, anti-CSD2 and anti-PrxQ (peroxiredoxin Q; both from Agrisera, Vännäs, Sweden) were used diluted 1:3000 and 1:1000 respectively, in Tris-buffered saline containing 0,1% Tween-20. The band optical densities were quantified using Image J. Analyses were performed in three biological replicates.

### TBARS assay

Lipid peroxidation was assayed using the TBARS (thiobarbituric acid reactive substances) assay as described in Larkindale and Knight (2002).

### Modelling of ACTIN2 protein structure

Samples of gDNA from *der1-3* mutant plants (from three different samples) were isolated using a phenol/chloroform/isoamylalcohol protocol (Pallotta *et al*., 2000). Isolated gDNA samples were subjected to sequencing (SeqMe, Czech Republic). Acquired sequences were compared with control *ACTIN2* gDNA sequence in Nucleotide BLAST database (https://blast.ncbi.nlm.nih.gov/Blast.cgi?PROGRAM=blastn&PAGE_TYPE=BlastSearch&LINK_LOC=blasthome). Sequences (both control and mutated) were translated to protein sequences in application http://bio.lundberg.gu.se/edu/translat.html?fbclid=IwAR3var5FJ8CBl4QqNe4Yic8NVz0TvWRd0TrF-uGUo6Nk6idLQxy2HvQqPEU. Only one single point mutation found (1114 C-T) changed protein sequence (Arg97-Cys97) accordingly. Protein sequences were used for protein structure modelling using application https://swissmodel.expasy.org/interactive?fbclid=IwAR1V9lhUgjiR1kUlwFLd8ojFftkHpkZwxIoT6mnEVIulEC2cPSYQov2twoE. The same application was used also for generation and downloading of representative images and videos.

### Data acquisition and analysis

Evaluated parameters such as root growth, skewness (representing an extent of actin filament bundling) and actin filament fluorescence integrated density (representing a percentage of occupancy) were measured in ImageJ (http://rsb.info.nih.gov/ij/). Graphs were prepared in Microsoft Excel program. Statistical significance between treatments at p < 0.05 was done using t-Test in Microsoft Excel or in program STATISTICA 12 (StatSoft) by ANOVA and subsequent Fisher’s LSD test (p<0,05).

## Results

### Impact of the der1-3 mutation and its topology on protein tertiary structure

*Arabidopsis thaliana* mutant *der1-3* (*deformed root hair1*) has been produced in the C24 ecotype background by an ethylmethanesulfonic acid - induced mutagenesis in the *DER1* locus, leading to a single-point mutation in the *ACTIN2* gene (Ringli *et al*., 2002). Mutants were selected according to a disturbed root hair development phenotype (Ringli *et al*., 2002, 2005). Single-point mutation in the gDNA sequence was determined at the position 1114 (changing cytosine to thymine), leading to altered protein sequence exchanging Arg97 to Cys97 in *der1-3* mutant (Ringli *et al*., 2002). We translated a nucleotide sequence, both in natural and mutated variant, to a primary protein sequence and we created a model of tertiary protein structure. We found that this position, both in natural (Fig. 1A) and mutated (Fig. 1B) ACTIN2 protein is placed in a loop located at the protein periphery (Suppl. Videos 1, 2). Importantly, the mutation does not alter the overall tertiary structure of protein (Fig. 1A, B), while this single amino acid exchange is topologically exposed to the protein surface (Fig. 1C, D). This analysis indicates that the ACTIN2 protein is produced in *der1-3* mutant and this single-point mutation exchanging Arg to Cys at the position 97 might rather influence its oxidation state, other protein post-translational modifications and/or protein-protein interactions.

**Figure 1.**
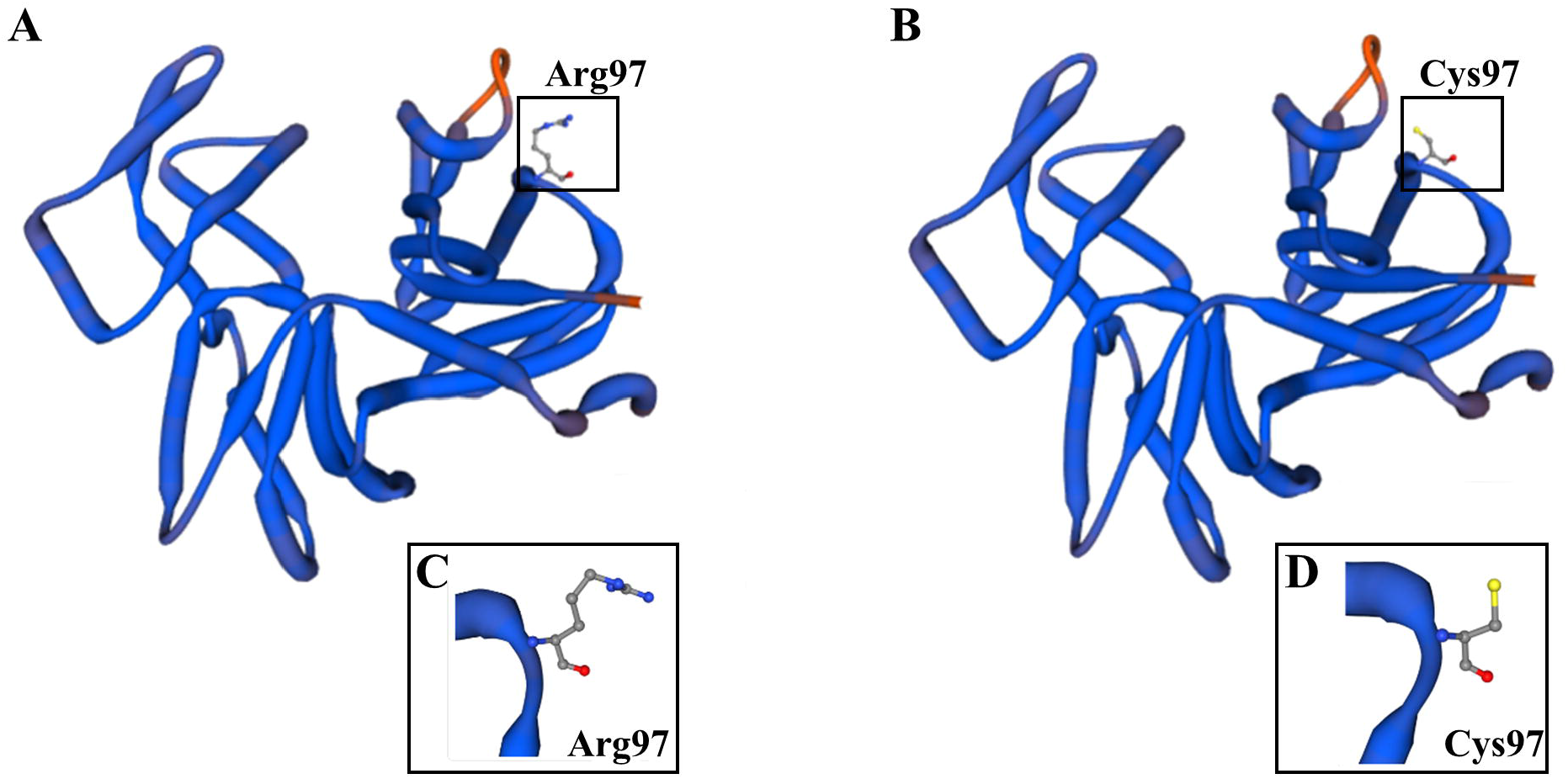
Model of the nature ACTIN2 protein structure and its mutated version in *der1-3* mutant. **(A-B)** SWISS model of the tertiary protein structure of ACTIN2 based on wild-type gene sequence **(A)** and based on gene sequence altered by single-point mutation in *der1-3* mutant **(B)**. Topological location of arginine in the position 97 of natural ACTIN2 **(A)** and substituted cysteine in the position 97 of mutated ACTIN2 **(B)** of *der1-3* mutant are showed in boxes. **(C-D)** Detailed structure of spatial arrangements of Arg97 **(C)** and Cys97 **(D)** from boxed area in **(A)** and **(B)**, respectively. 3D rotational models of protein structures are presented in Suppl. Movies 1 and 2. Models of protein structures were produced in: https://swissmodel.expasy.org/interactive?fbclid=IwAR1V9lhUgjiR1kUlwFLd8ojFftkHpkZwxIoT6mnEVIulEC2cPSYQov2twoE.

### Efficiency of seed germination under oxidative stress

In order to characterize responses of *der1-3* mutant to oxidative stress, we analysed several phenotypical parameters. Apart of obvious phenotype of root hairs that are arrested in tip growth after bulge formation (Ringli *et al*., 2002), mutant plants are affected in more developmental aspects. Among them, seeds of *der1-3* mutant germinate later than C24 wild-type seeds, and primary roots show more irregular and wavy growth pattern, due to a change in the cell division plane orientation (Vaškebová *et al*., 2018). Thus, our first interest was to test the rate of seed germination under conditions of mild and severe oxidative stress. Dry seeds of C24 and *der1-3* mutant after surface-sterilization and imbibition were tested for germination on solidified culture media with different concentrations of PQ. Germination efficiency of *der1-3* mutant seeds was lower than of C24 seeds under control conditions (Vaškebová *et al*., 2018), which was corroborated also in this study (Suppl. Fig. S1A). Treatment with three different concentrations of PQ (0.1, 0.2 and 0.5 μmol.l^-1^) did not influence considerably the rate of seed germination (Suppl. Fig. S1B-D). Thus, seed germination of *der1-3* was synchronously delayed comparing to C24 under PQ treatment (Suppl. Fig. S1B-D), similarly as in control conditions (Suppl. Fig. S1A).

The structure of actin cytoskeleton in cells of *der1-3* mutant was also compromised, showing thinner and less organized actin microfilaments in comparison to C24 control plants (Vaškebová *et al*., 2018). Based on this fact, we performed phenotypical analyses on transgenic C24 and *der1-3* mutant lines carrying a construct *pro35S::GFP:FABD2*, representing genetically-encoded marker for live cell imaging of actin cytoskeleton. Seed germination analysis revealed similar delay of germination efficiency in transgenic *der1-3* mutant line as compared to C24 transgenic line in both control conditions (Suppl. Fig. S1E) and after treatment with 0.1, 0.2 and 0.5 μmol.l^-1^ PQ (Suppl. Figs. S1F-H). Seed germination rate of transgenic C24 line carrying *pro35S::GFP:FABD2* (designated as C24 GFP-FABD2) and transgenic *der1-3* mutant line carrying *pro35S::GFP:FABD2* (designated as *der1-3* GFP-FABD2) within the first 24 h did not significantly differ from germination rate of C24 wild-type or *der1-3* mutant seeds, respectively, in control conditions (Suppl. Fig. S1I), as well as after treatment with 0.1, 0.2 and 0.5 μmol.l^-1^ PQ (Suppl. Fig. S1J-L). This experiment clearly showed that: I. seed germination is significantly delayed in *der1-3* mutant, II. expression of *pro35S::GFP:FABD2* construct in transgenic C24 and *der1-3* mutant lines did not affect the germination efficiency, and III. oxidative stress induced by PQ application did not affect considerably the physiological processes related to seed germination in both C24 and *der1-3* mutant.

### Influence of oxidative stress on post-germination root growth

Analysis of primary root growth of seedlings within the first 5 days after germination on control media revealed slightly lower elongation rate of *der1-3* roots in comparison to C24 (Fig. 2A), however, the differences in average root growth per 24 h were insignificant (Fig. 2I). We found the same root growth rate also in *der1-3* GFP-FABD2 line (Fig. 2I), but interestingly, average root growth rate per 24 h of C24 GFP-FABD2 line on control media was significantly higher (Fig. 2E, I). Thus, seedlings of transgenic C24 GFP-FABD2 line showed more effective root growth rate than C24 wild-type seedlings (Suppl. Fig. S2A), while there were no differences in this parameter between seedlings of *der1-3* and transgenic *der1-3* GFP-FABD2 line in control conditions (Suppl. Fig. S2B).

**Figure 2.**
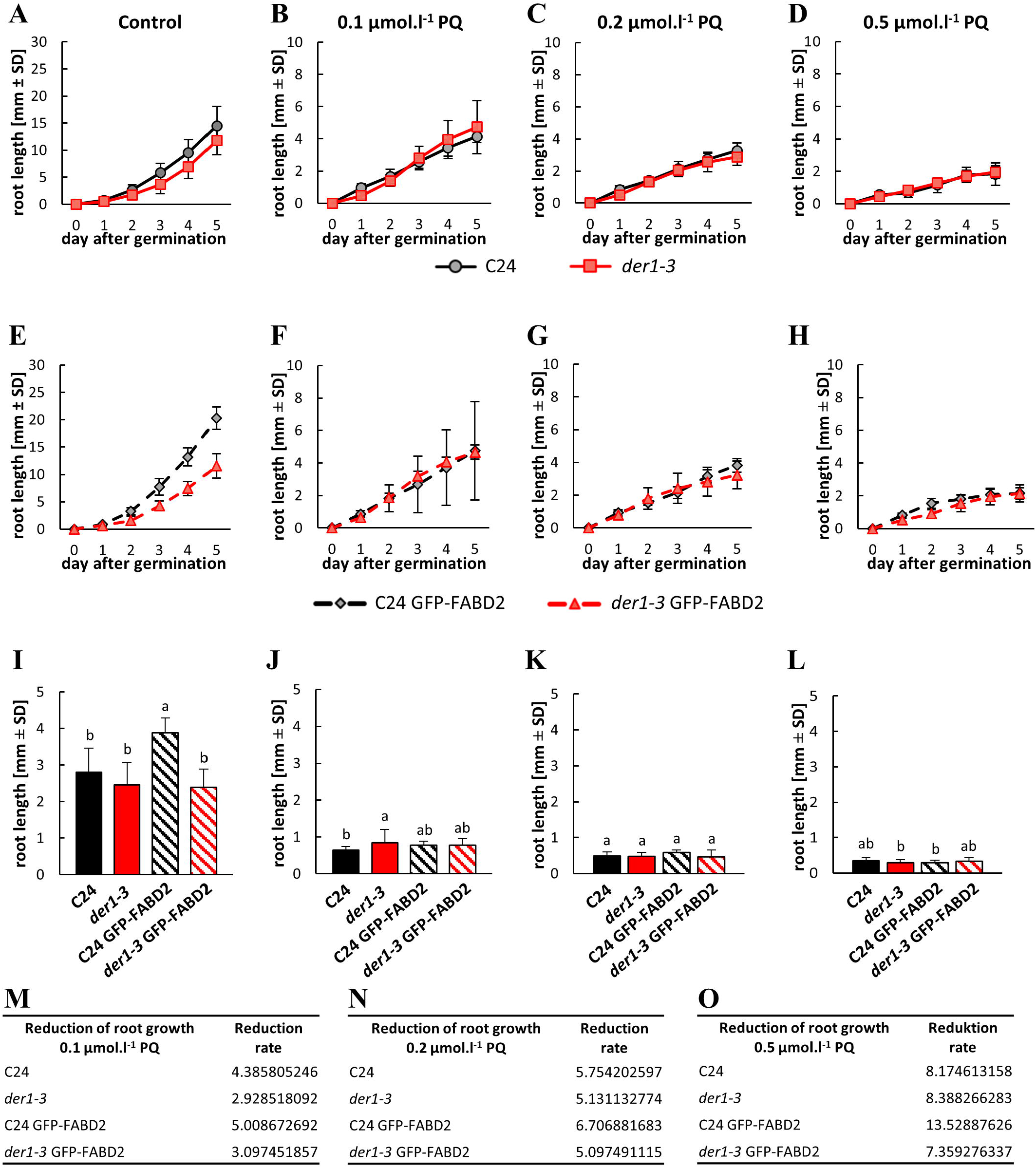
Root growth rate in plants of control C24, *der1-3* mutant and transgenic C24 and *der1-3* lines expressing *pro35S::GFP:FABD2* after germination in PQ-containing media. **(A-D)** Root growth rate within the first 5 days after germination of control C24 and *der1-3* mutant plants on control media **(A)** and on media containing 0.1 **(B)**, 0.2 **(C)** and 0.5 **(D)** μmol.l^-1^ PQ. **(E-H)** Root growth rate within the first 5 days after germination of transgenic C24 line carrying GFP-FABD2 and transgenic *der1-3* line carrying GFP-FABD2 on control media **(E)** and on media containing 0.1 **(F)**, 0.2 **(G)** and 0.5 **(H)** μmol.l^-1^ PQ. **(I-L)** Average root growth per 24 h on the control media **(I)** and on media containing 0.1 **(J)**, 0.2 **(K)** and 0.5 **(L)** μmol.l^-1^ PQ. **(M-O)** Reduction ratio (fold change in respect to control) of averaged root growth in respective lines on media containing 0.1 **(M)**, 0.2 **(N)** and 0.5 **(O)** μmol.l^-1^ PQ. Experiments were repeated two times with 16 plants per line (control) and 12 plants per line (PQ). Different lowercase letters above the bars **(I-L)** represent statistical significance according to one-way ANOVA and subsequent LSD test at p value < 0.05.

Seedlings of all tested lines germinating and growing on PQ-containing media within the first 5 days after germination showed reduction in primary root growth, which was dependent on PQ concentration (Suppl. Fig. S2C-F). Together with flattening of the root growth rate curves, there was also apparent PQ dose-dependent unification of the root growth rate between C24 wild-type and *der1-3* mutant seedlings (Fig. 2B-D), and also between C24 GFP-FABD2 and *der1-3* GFP-FABD2 seedlings (Fig. 2F-H). Dose-dependent reduction in average root growth per 24 h was apparent in seedlings germinating and growing on media containing 0.1, 0.2 and 0.5 μmol.l^-1^ PQ (Fig. 2J-L; Suppl. Fig. S2C-F). Although the root growth rate of *der1-3* mutant was always similar or lower in comparison to C24 wild-type under control conditions (Fig. 2A, I; Suppl. Fig. S2A, B), roots of *der1-3* mutant germinating and growing in the presence of 0.1 μmol.l^-1^ PQ showed better growth than C24 wild-type (Fig. 2J). Average root growth rate per 24 h was considerably reduced on media containing 0.2 and 0.5 μmol.l^-1^ PQ (Fig. 2K, L) without any differences among all tested lines. However, when differences were evaluated as reduction ratio in respect to control values, fold change in averaged root growth rate on media containing 0.1 μmol.l^-1^ PQ was 4.4 and 5.0 in C24 wild-type and C24 GFP-FABD2, respectively, but only 2.9 and 3.1 in *der1-3* mutant and *der1-3* GFP-FABD2, respectively (Fig. 2M). Although fold change in averaged root growth rate between C24 and *der1-3* mutant genotypes were less obvious on media containing 0.2 and 0.5 μmol.l^-1^ PQ, the reduction rate was similar or slightly lower in *der1-3* mutant (Fig. 2N, O). These data clearly indicate that root growth of both *der1-3* mutant and *der1-3* GFP-FABD2 transgenic line is less affected by mild and severe oxidative stress induced by PQ presence in the culture medium.

Effectivity of the root growth rate under oxidative stress in analysed lines was determined also by measurement of the distance between the first root hair and the root tip. In roots of 5 days-old plants growing in control conditions, this distance was significantly longer in C24 wild-type in comparison to *der1-3* (Suppl. Fig. S3A). The same tendency showing significantly longer distance between the first root hair and the root tip was observed also in transgenic C24 GFP-FABD2 line in comparison to transgenic *der1-3* GFP-FABD2 line (Suppl. Fig. S3A). Interestingly, both transgenic lines (C24 GFP-FABD2 and *der1-3* GFP-FABD2) had this measured distance significantly longer in comparison to control C24 and *der1-3* plants, respectively (Suppl. Fig. S3A). Such differences between C24 and *der1-3* mutant were reduced considerably or disappeared completely in seedlings germinating and growing on media containing 0.1, 0.2 and 0.5 μmol.l^-1^ PQ (Suppl. Figs. S3B-D). This represented another indication of differential responses to oxidative stress of analysed lines, showing significantly higher tolerance of transgenic *der1-3* mutant lines.

Plants monitored for 11 days after germination showed apparent time-dependent acceleration of root growth in control conditions (Suppl. Fig. S4A) with no obvious differences between C24 wild-type and *der1-3* mutant (Suppl. Fig. S4E). However, root growth of C24 GFP-FABD2 line was faster, particularly in later stages of development (Suppl. Fig. S4A), leading to significant increase of average root growth per 24 h (Suppl. Fig. S4E). Monitoring root growth rate upon prolonged PQ treatment showed clearly different trend of response between control C24 lines and *der1-3* mutant lines. On media containing 0.1 and 0.2 μmol.l^-1^ PQ, both *der1-3* mutant and *der1-3* GFP-FABD2 line showed better average root growth per 24 h as C24 wild-type and C24 GFP-FABD2 line (Suppl. Fig. S4F, G). Continuous monitoring of root growth rate revealed that it was higher in *der1-3* mutant and *der1-3* GFP-FABD2 line than in C24 wild-type and C24 GFP-FABD2 line from 8^th^ to 11^th^ day after germination (Suppl. Fig. S4B, C), and it was opposite to control conditions (Suppl. Fig. S4A). Root growth of plants on media containing 0.5 μmol.l^-1^ PQ was considerably reduced with minimal increase in length per day (Suppl. Fig. S4D), showing very similar average root growth per 24 h in all examined lines (Suppl. Fig. S4H). This analysis confirmed that, unlike the wild-type lines, root growth and development of *der1-3* mutant and *der1-3* GFP-FABD2 line is better adapted to the mild oxidative stress.

### Biomass production affected by oxidative stress

Shoot and root fresh weights analysed 18 days after germination of plants growing on control media revealed considerably higher biomass production in C24 wild-type and C24 GFP-FABD2 line. Oppositely, shoot and root biomass productions in *der1-3* mutant and *der1-3* GFP-FABD2 line were seemingly lower (Suppl. Fig. S5A). In plants germinating and 18 days growing on media containing PQ both shoot and root biomass production declined, but not uniformly among tested lines. While plants of C24 wild-type and C24 GFP-FABD2 line reacted to increasing concentrations of PQ by drastic reduction of both shoot and root biomass, it was not so dramatically reduced in *der1-3* mutant and *der1-3* GFP-FABD2 line. Already on media containing 0.1 and 0.2 μmol.l^-1^ PQ (Suppl. Figs. S5B, C) biomass weights of *der1-3* mutant and *der1-3* GFP-FABD2 line were similar or even higher as in C24 wild-type and C24 GFP-FABD2 line. Media with 0.5 μmol.l^-1^ PQ hindered massively root development, but shoot biomass production and development was clearly better in *der1-3* mutant and *der1-3* GFP-FABD2 line (Suppl. Fig. S5D). Calculation of the biomass production as a reduction ratio in fold change in respect to control revealed minimal change in both shoot and root biomass in *der1-3* mutant and *der1-3* GFP-FABD2 line on media containing 0.1 μmol.l^-1^ PQ in comparison to C24 wild-type and C24 GFP-FABD2 line (Suppl. Fig. S5E). On media containing 0.2 and 0.5 μmol.l^-1^ PQ (Suppl. Figs. S5F, G), we found higher biomass production (in fold change) in *der1-3* mutant and *der1-3* GFP-FABD2 line when compared to C24 wild-type and C24 GFP-FABD2 line. This analysis clearly revealed physiological resistance of *der1-3* mutant against mild and severe oxidative stress.

### Post-germination plant responses to oxidative stress

The process of seed germination in all tested lines was not considerably affected by oxidative stress induced by PQ presence in the culture medium (Suppl. Fig. S1). In order to characterize solely oxidative stress-related inhibition of root growth, we performed seed germination on control media and after that we transferred 3 days-old seedlings to culture media containing different concentrations of PQ. Comparison of root growth rate within 4 days after transfer showed that it is very similar for C24 wild-type and *der1-3* mutant on control media (Fig. 3A, I). After transfer of seedlings of transgenic lines, root growth rate on control media of C24 GFP-FABD2 line was significantly higher in comparison to *der1-3* GFP-FABD2 line (Fig. 3E). Actually, it was the highest among all tested lines (Fig. 3I). Root growth rate of C24 GFP-FABD2 line was higher as in C24 wild-type (Suppl. Fig. S6A), while there was no difference between *der1-3* mutant and *der1-3* GFP-FABD2 line (Suppl. Fig. S6B). Transfer of C24 wild-type and *der1-3* mutant seedlings germinated on control media to media containing 0.1, 0.2 and 0.5 μmol.l^-1^ PQ led to similarly decreased root growth rate (Fig. 3B, C, D). We found similar reaction also in seedlings of C24 GFP-FABD2 and *der1-3* GFP-FABD2 lines germinated on control media and transferred to media containing the same concentrations of PQ (Fig. 3F, G, H). Although the absolute root length of *der1-3* mutant and *der1-3* GFP-FABD2 line was lower on media containing 0.2 and 0.5 μmol.l^-1^ PQ (Fig. 3C, D, G, H), the averaged root growth rate was not considerably different (Fig. 3K, L). However, the reaction of seedlings to 0.1 μmol.l^-1^ PQ revealed much better tolerance of *der1-3* mutant and *der1-3* GFP-FABD2 line, as their averaged root growth rate was significantly higher than in C24 wild-type and C24 GFP-FABD2 line, respectively (Fig. 3J). Different mode of seedling reaction after transfer to media with 0.1 μmol.l^-1^ PQ was revealed. There was rather uniform reduction of the root growth rate on PQ-containing media in C24 wild-type and C24 GFP-FABD2 line (Suppl. Fig. S6C, D), while root growth rate much less affected by 0.1 μmol.l^-1^ PQ was clearly documented in *der1-3* mutant and *der1-3* GFP-FABD2 line (Suppl. Fig. S6E, F). These observations were corroborated by quantitative characterization of differences in averaged root growth rate by reduction ratio between control and PQ-containing media in fold changes. Root growth reduction ratio caused by 0.1 μmol.l^-1^ PQ was lower in seedlings of *der1-3* mutant and *der1-3* GFP-FABD2 line (Fig. 3M). Using 0.2 μmol.l^-1^ PQ, the differences between C24-related and *der1-3* mutant-related lines were lower in *der1-3* GFP-FABD2 line, while the reduction was stronger in *der1-3* mutant than in the C24 wild-type (Fig. 3N). Differences between lines transferred to 0.5 μmol.l^-1^ PQ were negligible (Fig. 3O).

**Figure 3.**
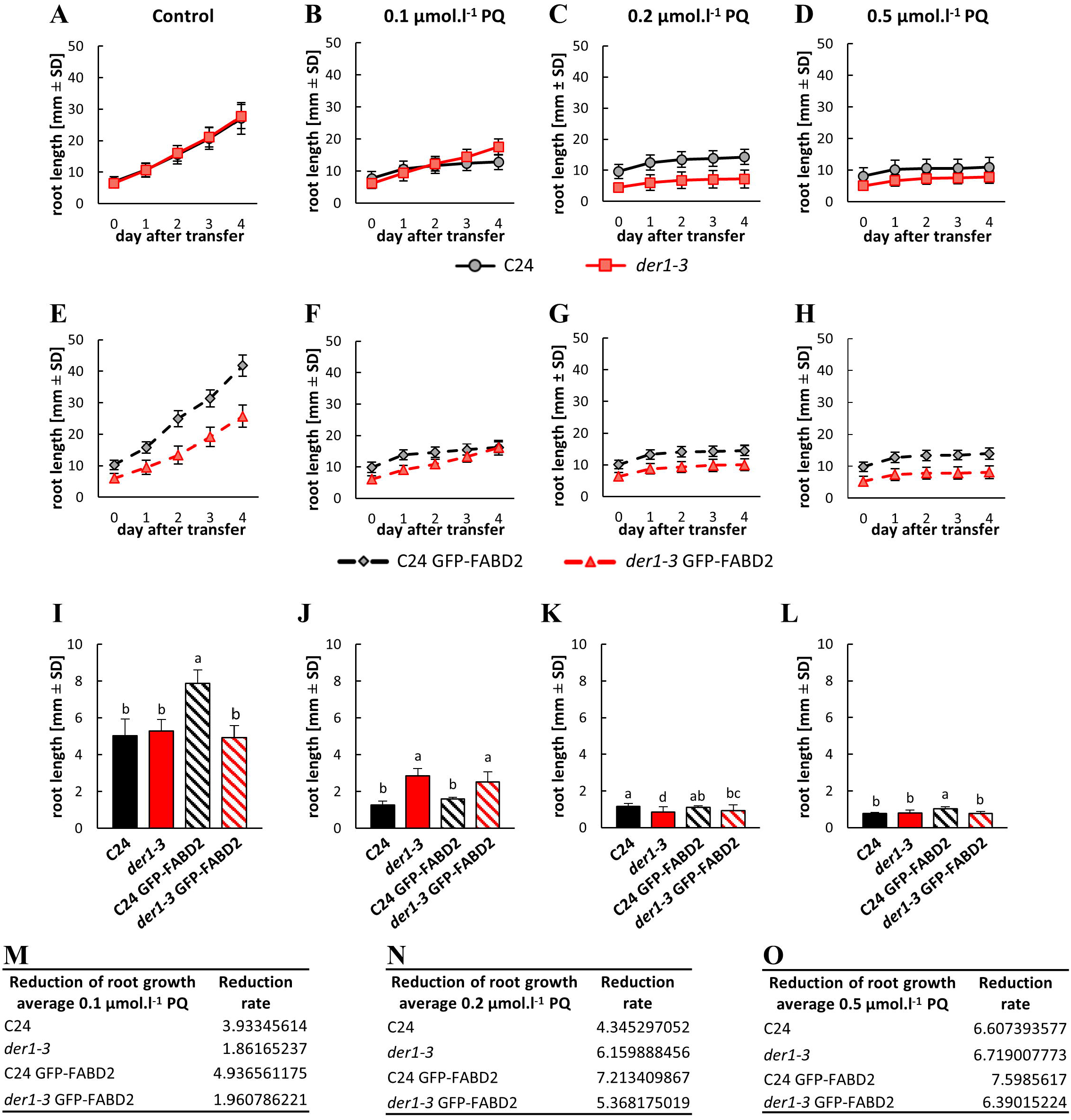
Root growth rate in plants of control C24, *der1-3* mutant and transgenic C24 and *der1-3* lines expressing *pro35S::GFP:FABD2* after their transfer to PQ-containing media. Plants 3 days old germinated on control media were transferred to PQ-containing media and root growth rate was analysed within subsequent 4 days. **(A-D)** Root growth rate of control C24 and *der1-3* mutant plants on control media **(A)** and on media containing 0.1 **(B)**, 0.2 **(C)** and 0.5 **(D)** μmol.l^-1^ PQ. **(E-H)** Root growth rate of transgenic C24 line carrying GFP-FABD2 and transgenic *der1-3* line carrying GFP-FABD2 on control media **(E)** and on media containing 0.1 **(F)**, 0.2 **(G)** and 0.5 **(H)** μmol.l^-1^ PQ. **(I-L)** Average root growth per 24 h on the control media **(I)** and on media containing 0.1 **(J)**, 0.2 **(K)** and 0.5 **(L)** μmol.l^-1^ PQ. **(M-O)** Reduction ratio (fold change in respect to control) of averaged root growth in respective lines on media containing 0.1 **(M)**, 0.2 **(N)** and 0.5 **(O)** μmol.l^-1^ PQ. Experiments were repeated two times with 16 plants per line (control) and 12 plants per line (PQ). Different lowercase letters above the bars **(I-L)** represent statistical significance according to one-way ANOVA and subsequent LSD test at p value < 0.05.

Phenotype of plants germinated and grown on control media for 20 days confirmed smaller above ground parts and more irregular and wavy root growth pattern of *der1-3* mutant (Suppl. Fig. S7A), in comparison to C24 wild-type (Suppl. Fig. S7B). Transgenic plants of C24 GFP-FABD2 and *der1-3* GFP-FABD2 grown on control media were bigger, but similar phenotypes, namely smaller above ground parts and more irregular and wavy root growth pattern of *der1-3* GFP-FABD2 line, were still apparent (Suppl. Fig. S7C, D). However, plants transferred from control to PQ-containing culture media revealed better development of *der1-3* mutant in comparison to C24 wild-type (Fig. 4A, B, C) and *der1-3* GFP-FABD2 line in comparison to C24 GFP-FABD2 line (Fig. 4D, E, F). In all concentrations of PQ tested, the above ground parts of *der1-3* mutant and *der1-3* GFP-FABD2 plants were much better developed (Fig. 4). In addition, considering purple pigmentation and arrested leaf enlargement, plants of *der1-3* mutant and *der1-3* GFP-FABD2 were much less affected in media containing 0.2 μmol.l^-1^ PQ (Fig. 4B, E) and 0.5 μmol.l^-1^ PQ (Fig. 4C, F). In media containing 0.1 μmol.l^-1^ PQ, plants of *der1-3* mutant and *der1-3* GFP-FABD2 line did not show such strong stress reaction and developmental arrest (Fig. 4A, D). Root development tested at the same conditions showed clear genotype-dependent response to PQ-induced oxidative stress. Plants (3 days-old) transferred from control to PQ-containing plates and photographed 17 days after transfer showed that *der1-3* mutant and *der1-3* GFP-FABD2 line, are much less sensitive to 0.1 μmol.l^-1^ PQ than C24 wild-type and C24 GFP-FABD2 line, respectively (Fig. 4A, D). Root system of *der1-3* mutant (Fig. 4A) and *der1-3* GFP-FABD2 line (Fig. 4D) maintained ability to grow and develop. Although 0.2 μmol.l^-1^ PQ considerably reduced root development, growing and branching capacity of *der1-3* mutant (Fig. 4B) and *der1-3* GFP-FABD2 line (Fig. 4E) were higher in comparison to C24 wild-type and C24 GFP-FABD2 line. Addition of 0.5 μmol.l^-1^ PQ to the culture medium reduced dramatically root development of all tested lines (Fig. 4C, F), which was apparent also from root growth rate (Fig. 3L) and root fresh weight (Suppl. Fig. S5D) analyses. Putting together, analyses of post-germination root growth and plant development after transfer to PQ-containing media from control conditions confirmed that plants of *der1-3* mutant and *der1-3* GFP-FABD2 line are more tolerant, particularly to the mild oxidative stress.

**Figure 4.**
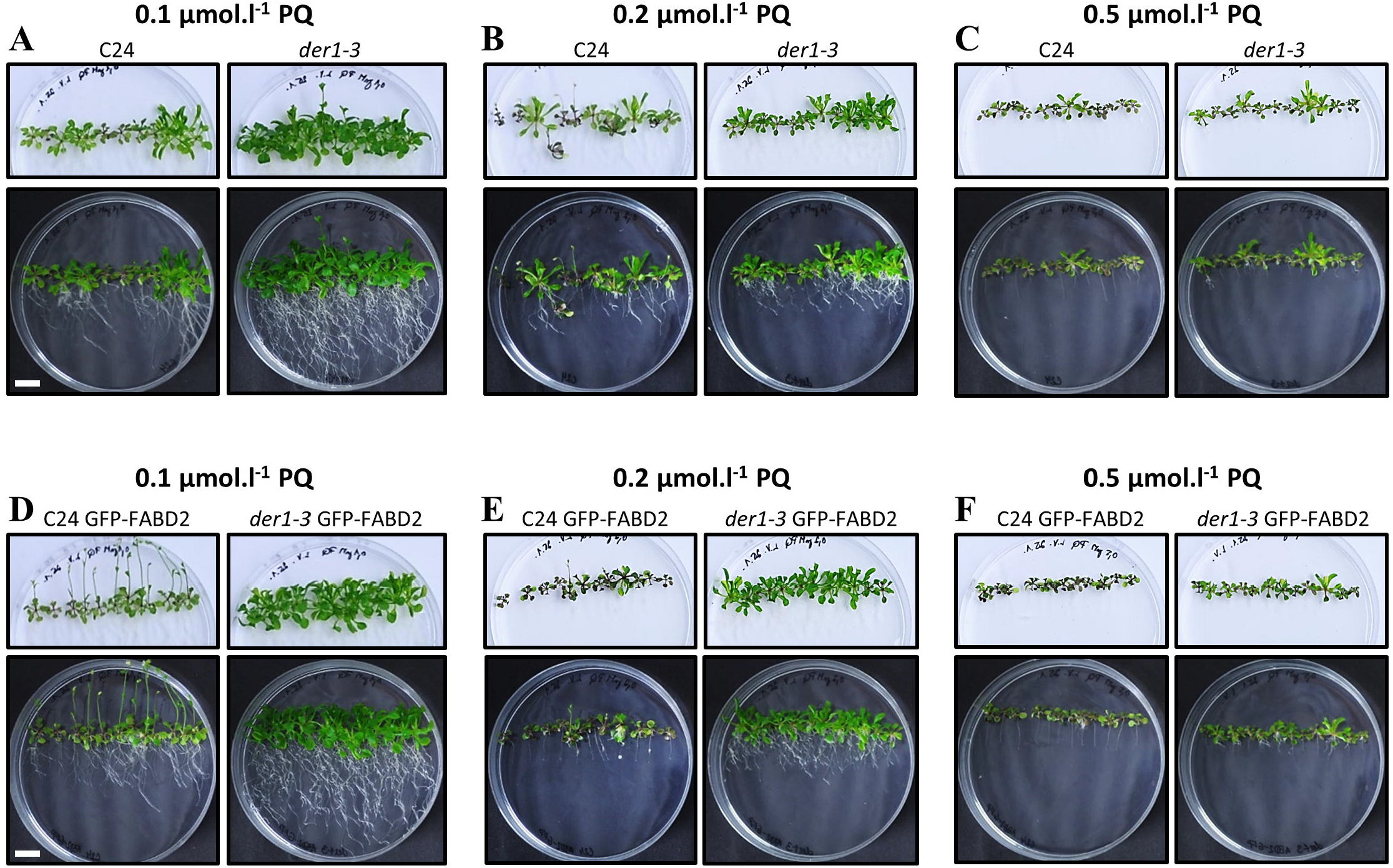
Plant phenotype of control C24, *der1-3* mutant and transgenic C24 and *der1-3* lines expressing *pro35S::GFP:FABD2* after their transfer to PQ-containing media. Plants 3 days old germinated on control media were transferred to PQ-containing media and photographed 17 days after transfer. **(A-C)** Plants of control C24 and *der1-3* mutant growing on media containing 0.1 **(A)**, 0.2 **(B)** and 0.5 **(C)** μmol.l^-1^ PQ. **(D-F)** Plants of transgenic C24 line carrying GFP-FABD2 and transgenic *der1-3* line carrying GFP-FABD2 growing on media containing 0.1 **(D)**, 0.2 **(E)** and 0.5 **(F)** μmol.l^-1^ PQ. Aboveground parts of plants were photographed on white background (upper row of images), and whole plants including roots were documented on black background (lower row of images). Plants grown on control media are documented in Suppl. Figure S7. Scale bar = 1 cm.

Taking into account the inhibitory effects of PQ in photosynthetically-active plant tissues, we employed also H_2_O_2_ treatment, as an alternative oxidative stress-inducing agent that directly affects the root system and its development. Four different concentrations of H_2_O_2_ (0.5; 1; 1.5 and 3 mmol.l^-1^) were tested in post-germination root growth rate analysis within 4 days after transfer of 3 days-old seedlings germinated on control media. We observed H_2_O_2_ dose-dependent response in the inhibition of root elongation (Fig. 5A-D). Averaged root length of both compared lines was gradually reduced by the presence of 0.5, 1, 1.5 and 3 mmol.l^-1^ H_2_O_2_ (Figs. 5A-E) in the culture medium. We observed significantly longer roots of C24 wild-type plants than *der1-3* mutant plants within the testing period in control conditions, and on media containing 0.5 and 1 mmol.l^-1^ H_2_O_2_, however, there was no statistically significant difference in the root length on media containing 1.5 mmol.l^-1^ H_2_O_2_ (Fig. 5E). Interestingly, the stronger concentration of H_2_O_2_ tested (1.5 mmol.l^-1^) inhibited root elongation in C24 wild-type significantly more than in *der1-3* mutant plants (Fig. 5E).

**Figure 5.**
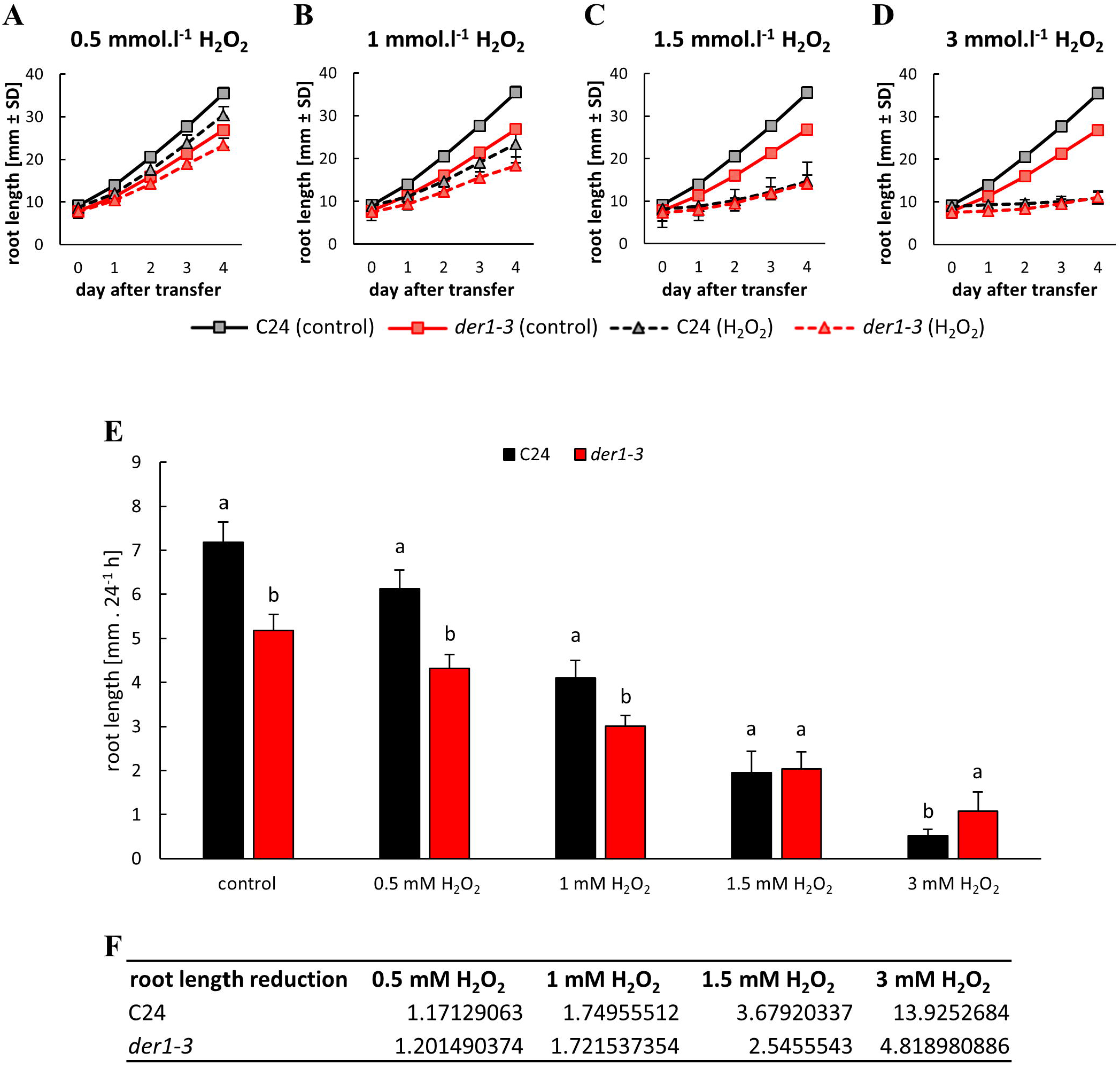
Root growth rate of control C24 and *der1-3* mutant plants after their transfer to H_2_O_2_-containing media. Plants 3 days old germinated on control media were transferred to H_2_O_2_-containing media and root growth rate was analysed within subsequent 4 days. **(A-D)** Root growth rate of control C24 and *der1-3* mutant plants on media containing 0.5 **(A)**, 1 **(B)**, 1.5 **(C)** and 3 **(D)** mmol.l^-1^ H_2_O_2_. **(E)** Average root growth per 24 h on the control media **(I)** and on media containing indicated concentrations of H_2_O_2_. **(F)** Reduction ratio (fold change in respect to control) of averaged root growth in control C24 and *der1-3* mutant plants on media containing 0.5 **(A)**, 1 **(B)**, 1.5 **(C)** and 3 **(D)** mmol.l^-1^ H_2_O_2_. Experiments were repeated two times with 10 plants per line. Different lowercase letters above the bars **(E)** represent statistical significance according to one-way ANOVA and subsequent LSD test at p value < 0.05.

Quantitative characterization of differences in averaged root growth rate presented as a reduction ratio between control and H_2_O_2_-containing media showed no differences between C24 wild-type and *der1-3* mutant on media containing 0.5 and 1 mmol.l^-1^ H_2_O_2_ (Fig. 5F). However, moderate difference caused by 1.5 mmol.l^-1^ H_2_O_2_ and considerably increased difference induced by 3 mmol.l^-1^ H_2_O_2_ (Fig. 5F) suggested that root growth and development of *der1-3* mutant plants are substantially more resistant to moderate and severe oxidative stress than of C24 wild-type plants. It can be further documented also by phenotype of whole plants. Together with the root system that was severely reduced by increasing concentration of H_2_O_2_ in C24 wild-type plants, reduction in the development of their above ground parts was also obvious (Suppl. Fig. S8A). In comparison, although the development of root system of *der1-3* mutant plants was also accordingly reduced by increasing concentration of H_2_O_2_, development of their above ground parts was less affected (Suppl. Fig. S8B). The overall data of phenotypical analyses thus indicate that *der1-3* mutant and transgenic plants in *der1-3* mutant background maintain growth and development because they are better protected against PQ- or H_2_O_2_-induced oxidative stress.

### Oxidative stress and response of the actin cytoskeleton

In order to characterize organization and dynamic properties of actin cytoskeleton under PQ-induced oxidative stress, we utilized transgenic C24 and *der1-3* lines expressing *pro35S::GFP:FABD2* construct. In hypocotyl epidermal cells of 3 days-old plants of C24 GFP-FABD2 line in control conditions, actin filaments were arranged in extensive, well organized and dynamic network (Suppl. Video 3). However, we observed massive bundling, particularly in cortical layers of the cell after treatment with 0.1 μmol.l^-1^ PQ for 30 min (Fig. 6A; Suppl. Video 4). Semi-quantitative evaluation of actin filament skewness, determining a degree of actin filaments bundling, showed increased levels after application of oxidative stress (Fig. 6B). Semi-quantitative evaluation of integrated density, determining fluorescence signal intensity per 1 μm^2^, was also significantly increased after PQ treatment (Fig. 6C). Actin filament organization was slightly different in hypocotyl epidermal cells of 3 days-old *der1-3* GFP-FABD2 plants in control conditions, showing mainly thinner, less organized, but dynamic actin filaments in cell cortex (Suppl. Video 5). Treatment with 0.1 μmol.l^-1^ PQ for 30 min induced partial bundling of actin filaments, but overall changes in their organization and dynamics were not so dramatic (Fig. 6D; Suppl. Video 6). As a result, both actin filament skewness, determining a degree of actin filaments bundling (Fig. 6E), and integrated density, determining mean fluorescence signal intensity (Fig. 6F), were not significantly affected by oxidative stress.

**Figure 6.**
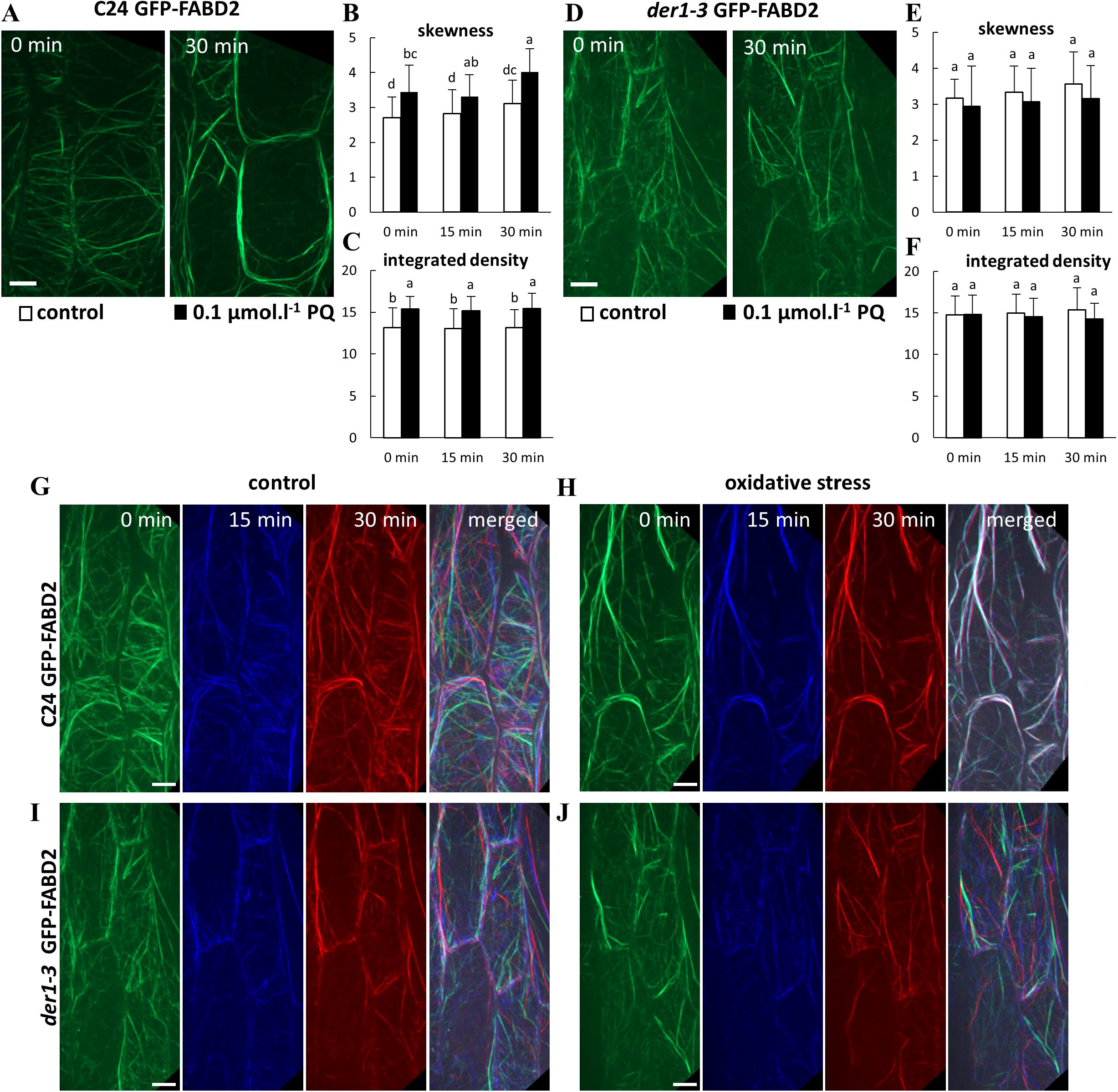
Organization and dynamics of actin filaments in hypocotyl epidermal cells of transgenic C24 and *der1-3* lines expressing *pro35S::GFP:FABD2* under PQ-induced oxidative stress. **(A-C)** Actin filaments in hypocotyl epidermal cells of 3 days-old plant of C24 GFP-FABD2 line in control conditions and after treatment with 0.1 μmol.l^-1^ PQ for 30 min **(A)**. Quantitative analysis of actin filaments bundling extent (skewness, **B**) and actin filaments density (percentage of occupancy, **C**) in control conditions and after application of 0.1 μmol.l^-1^ PQ. **(D-F)** Actin filaments in hypocotyl epidermal cells of 3 days-old plant of *der1-3* GFP-FABD2 line in control conditions and after treatment with 0.1 μmol.l^-1^ PQ for 30 min **(D)**. Quantitative analysis of actin filaments bundling extent (skewness, **E**) and actin filaments density (percentage of occupancy, **F**) in control conditions and after application of 0.1 μmol.l^-1^ PQ. Data were analysed on images collected from hypocotyl epidermal cells within 0, 15 and 30 min time-points of scanning. (**G-J**) Semiquantitative analysis of actin filament dynamics in hypocotyl epidermal cells presented by pseudocolouring displacement analysis. Dynamic properties of actin filaments in C24 GFP-FABD2 line in control conditions **(G)** and after application of 0.1 μmol.l^-1^ PQ **(H)**. Dynamic properties of actin filaments in *der1-3* GFP-FABD2 line in control conditions **(I)** and after application of 0.1 μmol.l^-1^ PQ **(J)**. Images acquired at the beginning, after 15 min and after 30 min of time-point scanning were coloured red, green, and blue, respectively, and merged. White colour indicates lowering (eventually stopping) of the actin dynamic activity. Experiments were repeated 5-6 times with 4-5 cells per plant in each line. Different lowercase letters above the bars **(B, C, E, F)** represent statistical significance according to one-way ANOVA and subsequent LSD test at p value < 0.05. Scale bar = 10 μm.

Next, dynamic properties of actin cytoskeleton in hypocotyl epidermal cells were analysed by sequential imaging of actin filaments within 30 min (by acquiring 0, 15 and 30 min time-points), followed by pseudocolour–based evaluation of their lateral displacement. In control conditions, dynamic changes of actin filament network in cells of both C24 GFP-FABD2 (Fig. 6G; Suppl. Video 3) and *der1-3* GFP-FABD2 (Fig. 6I; Suppl. Video 5) lines were determined by minimal overlay of sequential coloured scans in merged images. After application of 0.1 μmol.l^-1^ PQ the same analysis revealed formation of excessive actin bundles with minimal dynamic changes in structure and organization in cells of C24 GFP-FABD2 line (Fig. 6H; Suppl. Video 4), while much less bundles were formed in cells of *der1-3* GFP-FABD2 line. In addition, overlay of sequential coloured scans revealed unchanged dynamic properties that was still high particularly in fine actin filaments (Fig. 6J; Suppl. Video 6). This analysis showing alterations in structure and dynamic properties of actin cytoskeleton in *der1-3* GFP-FABD2 line, that were not considerably affected by PQ treatment, may significantly support observed physiological resistance of *der1-3* mutant and related transgenic *der1-3* GFP-FABD2 line against oxidative stress.

### ROS production, lipid peroxidation and antioxidant activity

Relative levels of ROS were determined by the histochemical detection of their production in cotyledons and leaves of seedlings under control and oxidative stress-inducing conditions. Visualization was performed using NBT staining of O2^•−^ production and DAB staining of H_2_O_2_ production, respectively. Semi-quantitative evaluation of the mean staining intensity revealed that both in cotyledons (Suppl. Fig. S9A) and leaves (Suppl. Fig. S9B) of plants treated for 7 days was no difference in O2^•−^ production upon PQ and H_2_O_2_ treatments between C24 wild-type and *der1-3* mutant. The ability of H_2_O_2_ production visualised by DAB staining was lower in *der1-3* mutant than in C24 wild-type in control conditions, both in cotyledons (Suppl. Fig. S9C) and leaves (Suppl. Fig. S9D). Interestingly, upon PQ and H_2_O_2_ treatments, the level of H_2_O_2_ production in *der1-3* mutant was increased to the C24 wild-type level, both in cotyledons and leaves (Suppl. Fig. S9C, D).

Based on observed phenotypical differences, we aimed to provide an evidence about the biochemical mechanisms underlying an increased tolerance of *der1-3* mutant to oxidative stress. Our analyses showed that *der1-3* mutant exhibited lower degree of lipid peroxidation after long-term PQ treatment compared to C24 wild-type, while cultivation on H_2_O_2_-containing media did not cause lipid peroxidation in any examined line (Fig. 7A). This indicated that PQ treatment was less damaging to *der1-3* mutant as compared to C24 in terms of membrane integrity. Next, we also examined activities of important antioxidant enzymes in *der1-3* mutant and C24 wild-type. We have found elevated capacity to decompose superoxide in *der1-3* mutant, as manifested by more intensive activation of iron superoxide dismutase 1 (FeSOD1) in these plants when exposed to both PQ and H_2_O_2_, as compared to C24 wild-type (Fig. 7B, C). Both treatments substantially decreased the activity of copper-zinc superoxide dismutase (CuZnSOD) isoforms in both *der1-3* mutant and C24 wild-type, while this reduction was less pronounced in *der1-3* mutant (Fig. 7B, C). This was observed also on protein abundance levels, as shown by immunoblotting (Fig. 7D, E). In addition, we encountered also substantially increased abundance of chloroplastic PrxQ, a H_2_O_2_ decomposing enzyme, in *der1-3* mutant in response to both PQ and H_2_O_2_ treatments, while slight increase (in the case of PQ), or unchanged abundance (in the case of H_2_O_2_), were observed in C24 wild-type (Fig. 7F, G). Altogether, these results showed that increased tolerance of *der1-3* mutant plants to oxidative stress is determined by elevated enzymatic capacity to decompose reactive oxygen species.

**Figure 7.**
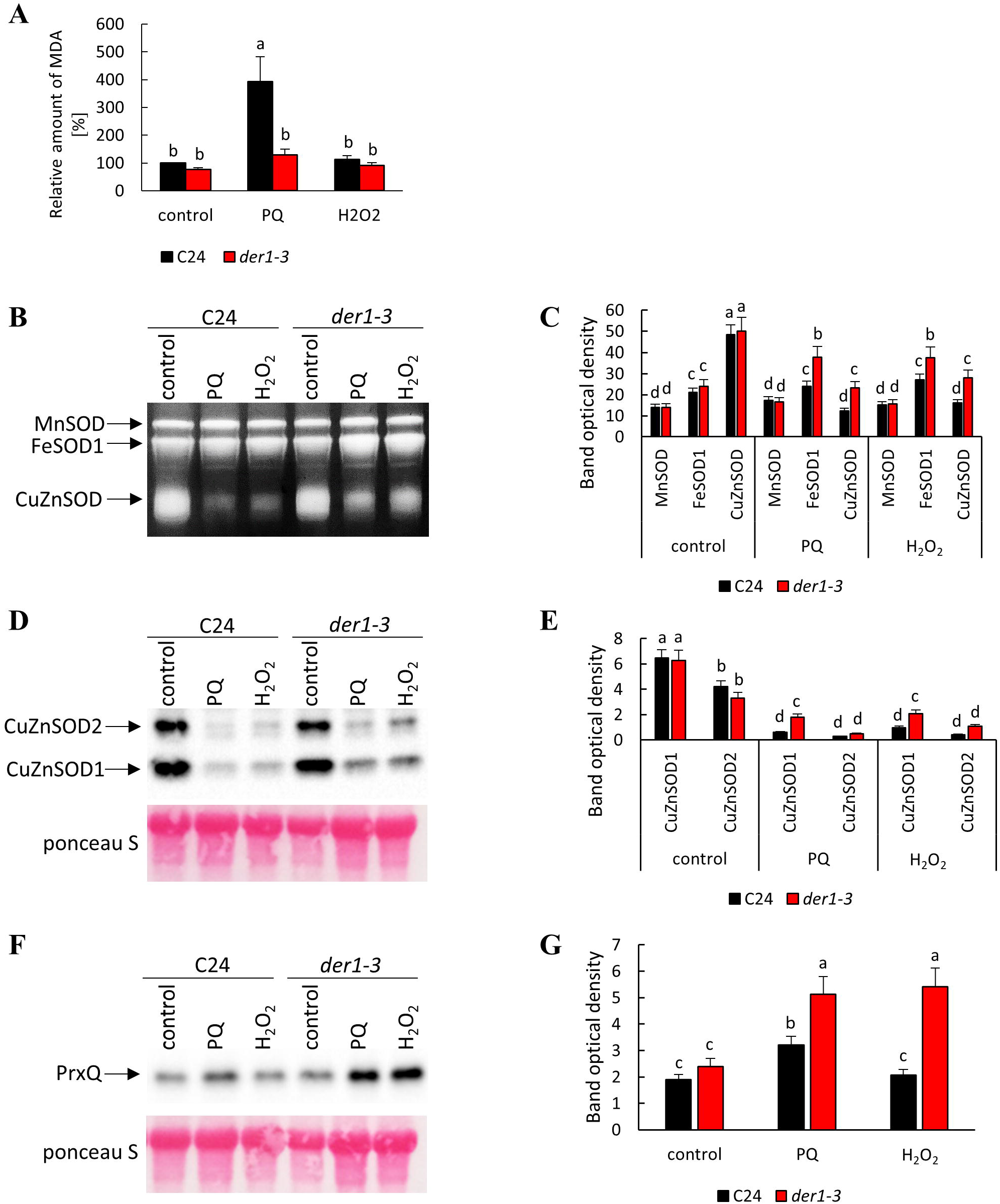
Estimation of lipid peroxidation and antioxidant capacity in plants of C24 wild-type and *der1-3* mutant. Plants 3 days old germinated on control media were transferred to 0.1 μmol.l^-1^ PQ- and 3 mmol.l^-1^ H_2_O_2_-containing media. **(A)** Relative quantification of malondyaldehyde content. **(B-C)** Visualisation of superoxide dismutase (SOD) isoforms activity **(B)** and quantification of individual band densities **(C)** on native polyacrylamide gels. **(D-E)** Immunoblot of CuZnSOD1 and CuZnSOD2 isoforms **(D)** and quantification of band densities **(E)**. **(F-G)** Immunoblot of peroxiredoxin Q abundance **(F)** and quantification of band densities **(G)**. Different lowercase letters above the bars **(A, C, E, G)** represent statistical significance between treatments according to t-Test at p value < 0.05.

## Discussion

Random mutagenesis approach in *Arabidopsis thaliana* led to the isolation of *der1* mutants, bearing a single-point mutation in *ACTIN2* gene (Ringli *et al*., 2002). It was demonstrated that these mutants (*der1-1, der1-2, der1-3*) are compromised in root hair development after the bulge initiation, showing typical “short root hair” phenotypes. Locations of these mutations to the *ACTIN2* gene confirmed essential role of actin cytoskeleton in the root hair development (Ringli *et al*., 2002, 2005). Further phenotypical, developmental and microscopic analyses of *der1-3* mutant plants revealed that seeds of *der1-3* mutant germinated later than those of C24 wild-type (Vaškebová *et al*., 2018). Moreover, due to changes in the cell division plane orientation, primary roots showed irregular and wavy pattern while actin filaments in epidermal cells of different plant organs (roots, hypocotyls and cotyledons) were shorter, thinner and arranged in more random orientations in the *der1-3* mutant (Vaškebová *et al*., 2018). Thus, *der1-3* mutant was affected in broader range of morphological and developmental aspects, related to alterations of the actin cytoskeleton and its organization at cellular level. It is not clear how structural and dynamic properties of actin cytoskeleton may support plant reactions to oxidative stress, therefore we addressed this in the present study. We induced conditions of mild and severe oxidative stress by supplementing PQ or H_2_O_2_ to the culture medium and characterized diverse parameters like germination, growth, development and biomass production in *der1-3* mutant and C24 wild-type plants. Analyses were done on plants germinating directly on such oxidative stress-inducing media or on seedlings germinated on control media first and subsequently transferred to PQ- or H_2_O_2_-containing plates.

Experiments on seed germination corroborated previously published data about slightly later germination of *der1-3* mutant (Vaškebová *et al*., 2018). We found that oxidative stress induced by PQ in concentrations, which has clear negative effect on root growth, did not affect the processes of seed germination in both C24 plants and *der1-3* mutant. Nevertheless, PQ in higher concentrations negatively affects seed germination (Haslekås *et al*., 2003) indicating that processes connected to germination are more resistant to PQ compared to root growth. Next, we prepared transgenic lines of C24 and *der1-3* expressing *pro35S::GFP:FABD2* construct in order to perform live-cell microscopic characterization of actin cytoskeleton, its organization and dynamic properties in these lines. We also characterized root growth rate in all above-mentioned lines. Interestingly, root growth of both *der1-3* mutant and *der1-3* GFP-FABD2 transgenic line was much less affected by mild and severe PQ-induced oxidative stress as in control C24, particularly at concentration 0.1 μmol.l^-1^ PQ in the culture medium. As a consequence, the reduction ratio in average root growth quantified as a fold change in respect to the C24 control was several orders lower in *der1-3* mutant (Fig. 2M, 3M, 5F). We found a similar tendency in the reduction of biomass production by PQ treatment, which was several orders stronger in C24 wild-type than in *der1-3* mutant. This trend was documented in both root and shoot biomass production and recorded in all PQ concentrations tested (Suppl. Fig. S5E, F, G). All data from analyses of post-germination root growth and plant development, both germinating on PQ-containing media and after transfer of non-treated seedlings to PQ-containing media, indicate that plants of *der1-3* mutant and *der1-3* GFP-FABD2 line are more tolerant, particularly to the mild oxidative stress.

Observed changes of phenotypical parameters distinguishing C24 wild-type from *der1-3* mutant indicate different sensitivity to oxidative stress. In addition, we performed several supporting biochemical experiments. Estimation of lipid peroxidation based on relative quantification of malondyaldehyde content revealed that *der1-3* mutant exhibits lower degree of lipid peroxidation after long-term PQ treatment compared to C24 wild-type. Thus, an important aspect of antioxidant defence in plants, namely membrane integrity, was better protected in *der1-3* mutant. Better tolerance of *der1-3* mutant against oxidative stress was supported also by abundance and activity of antioxidant enzymes such as iron superoxide dismutase 1 (FeSOD1) and two copper-zinc superoxide dismutase isoforms (CuZnSOD1 and CuZnSOD2). Elevated levels of FeSOD1 (activity) and CuZnSOD1/2 (activity and abundance) in *der1-3* mutant after long-term PQ and H_2_O_2_ exposure point to higher capacity of the mutant to decompose superoxide radical compared to C24 wild-type. CuZnSOD1/2 and FeSOD1 are proposed as important determinants of oxidative stress tolerance (Sunkar *et al*., 2006; Dvořák *et al*., 2020). *der1-3* mutants possess also increased H_2_O_2_ decomposing efficiency which is executed by PrxQ. Nevertheless, other mechanisms of H_2_O_2_ removal cannot be excluded. PrxQ is an atypical 2-cys peroxiredoxin which uses (and interacts with) thioredoxin as an electron donor to decompose H_2_O_2_ (Lamkemeyer *et al*., 2006; Yoshida *et al*., 2015). Peroxiredoxins and thioredoxins as redox buffering proteins, in addition, may modulate intracellular signalling related to ROS (Dietz *et al*., 2006). Thus, we propose that higher capacity to decompose ROS and enhanced cellular redox regulation might be the main factors determining an increased tolerance of *der1-3* mutant to oxidative stress.

The next task was to reveal how structure and organization of actin cytoskeleton in *der1-3* may support increased tolerance of this mutant to oxidative stress. Previous study reported that *der1-3* mutant does not show solely root hair phenotype, but the actin cytoskeleton was altered and affected also root growth and development. Actin filaments in cells of *der1-3* mutant were shorter, thinner and arranged in more random orientations (Vaškebová *et al*., 2018). Oxidative stress caused by application of 0.1 μmol.l^-1^ PQ for 30 min induced massive bundling of actin filaments in cells of C24 GFP-FABD2 line. Actin cytoskeleton in cells of *der1-3* GFP-FABD2 line was arranged in extensive network, although actin filaments were thinner and less organized. However, this organization was virtually insensitive to 0.1 μmol.l^-1^ PQ applied for 30 min. A higher protection of actin filaments fine network in *der1-3* GFP-FABD2 line was accompanied by roughly unchanged dynamic properties under PQ treatment. Thus, higher resistance of the actin cytoskeleton against deteriorating effects of oxidative stress may be one of the main molecular mechanism supporting the higher tolerance of *der1-3* mutant to this type of stress. This can be related to proposed role of actin cytoskeleton in adaptation of Arabidopsis root meristem cells to oxidative stress through protecting PIN2 auxin efflux carrier trafficking to the plasma membrane, which is controlled by auxin levels. Since auxin levels were disturbed by generated ROS, the abundance of PIN2 at the plasma membrane decreased. The role of actin cytoskeleton lies on keeping the PIN2 intracellular trafficking, which requires the function of the ADP-ribosylation factor (ARF)-guaninenucleotide exchange factor (GEF) BEN1, an actin-associated regulator (Zwiewka *et al*., 2019). However, it is not known how this and similar functions can be affected by altered structural and dynamic properties of the actin cytoskeleton in *der1-3* mutant. It was proposed that PIN2 intracellular trafficking was reduced because H_2_O_2_ treatment affected actin dynamics (Zwiewka *et al*., 2019). Reduction in actin filament bundling can be directly associated with increased actin filament dynamics (Staiger and Blanchoin, 2006). Similarly, treatment of Arabidopsis plants with strigolactones reduces bundling of actin filaments with their simultaneously increasing dynamics, however, *der1-2* and *der1-3* mutants were much less sensitive to strigolactone analogue GR24 (Pandya-Kumar *et al*., 2014). Collectively, these data support our conclusion that actin filament arrangement less prone to bundling and staying dynamic is critical for actin properties in *der1-3* mutant, significantly contributing also to higher tolerance of this mutant against oxidative stress.

Position of the mutated amino acid in the ACTIN2 sequence, Arg-97, is located in the subdomain 1 on the protein surface (Fig. 1A; Suppl. Movie 1; Diet *et al*., 2004; Kabsch *et al*., 1990; Ringli *et al*., 2002). In mutated ACTIN2, *der1-3* mutation causes the single amino acid exchange of Arg by Cys, which is topologically exposed to the protein surface. Although it probably does not influence the protein tertiary structure, the topology of this modification might have strong impact on the post-translational modifications of ACTIN2, or its ability to perform protein-protein interactions in *der1-3* mutant. There are some supporting facts for this. BEN1, a guanine exchange factor for ARF, regulating actin filament-based intracellular trafficking of PIN2 during adaptation to oxidative stress, contains highly conserved cysteine residues (Mouratou *et al*., 2005; Zwiewka *et al*., 2019) that could be modified by H_2_O_2_ treatment. Increased redox status upon accumulation of H_2_O_2_ can initiate oxidation of cysteine sulfhydryl groups in actins (Wang *et al*., 2012). As mutated ACTIN2 protein in *der1-3* mutant contains additional Cys compared to the native one, we hypothesize that ACTIN2 in *der1-3* might undergo redox-mediated posttranslational modifications accelerating, via PrxQ and thioredoxins, the antioxidant capacity in *der1-3* mutant.

Putting together, our data indicate that topologically important change in ACTIN2 in the *der1-3* mutant is linked to the better tolerance to mild and severe oxidative stress, increased capacity to decompose ROS and higher dynamicity of the actin cytoskeleton.

## Supplementary Information

*Suppl. Fig. S1*. Seed germination under the PQ treatment of control C24, *der1-3* mutant and transgenic C24 and *der1-3* lines expressing *pro35S::GFP:FABD2*.

*Suppl. Fig. S2*. Impact of PQ treatment on root growth rate in control and transgenic C24 and *der1-3* mutant lines.

*Suppl. Fig. S3*. Effect of PQ treatment on the distance between the first root hair and the root tip in control and transgenic C24 and *der1-3* mutant lines.

*Suppl. Fig. S4*. Root growth rate in plants of control C24, *der1-3* mutant and transgenic C24 and *der1-3* lines under prolonged PQ treatment.

*Suppl. Fig. S5*. Shoot and root fresh weight in plants of control C24, *der1-3* mutant and transgenic C24 and *der1-3* lines expressing *pro35S::GFP:FABD2* after germination and growth in PQ-containing media.

*Suppl. Fig. S6*. Root growth rate in control and transgenic C24 and *der1-3* mutant lines after their transfer to PQ-containing media

*Suppl. Fig. S7*. Plant phenotype of control and transgenic C24 and *der1-3* mutant lines on control media.

*Suppl. Fig. S8*. Phenotype of control C24 and *der1-3* mutant plants after their transfer to H_2_O_2_-containing media.

*Suppl. Fig. S9*. Histochemical detection of O2^•−^ and H_2_O_2_ production in cotyledons and leaves of control C24 and *der1-3* mutant plants after their transfer to PQ- and H_2_O_2_-containing media.

*Suppl. Video 1*. 3D rotational model of protein structure of the nature ACTIN2 protein.

*Suppl. Video 2*. 3D rotational model of protein structure of mutated version in *der1-3* mutant.

*Suppl. Video 3*. Actin filaments in hypocotyl epidermal cells of C24 GFP-FABD2 line recorded in 30 seconds intervals for 30 min in control conditions.

*Suppl. Video 4*. Actin filaments in hypocotyl epidermal cells of C24 GFP-FABD2 line recorded in 30 seconds intervals after treatment with 0.1 μmol.l^-1^ PQ for 30 min.

*Suppl. Video 5*. Actin filaments in hypocotyl epidermal cells of *der1-3* GFP-FABD2 line recorded in 30 seconds intervals for 30 min in control conditions.

*Suppl. Video 6*. Actin filaments in hypocotyl epidermal cells of *der1-3* GFP-FABD2 line recorded in 30 seconds intervals after treatment with 0.1 μmol.l^-1^ PQ for 30 min.

## Acknowledgements

The work was supported by Czech Science Foundation GAČR, project Nr. 19-18675S and by ERDF project “Plants as a tool for sustainable global development” (No. CZ.02.1.01/0.0/0.0/16_019/0000827).

## Author contributions

L.K. performed phenotypical and microscopic analyses, data processing and statistical evaluation. T.T. performed biochemical analyses. M.O. and L.K. wrote the manuscript with input from all co-authors. M.O. and J.Š. conceived the study, designed the experiments and made final editing. J.Š. provided infrastructure, coordinated the whole project and supervised the work.

## Data availability statement

All data supporting the findings of this study are available within the paper and within its supplementary materials published online.

## References

Baluška F, Salaj J, Mathur J, Braun M, Jasper F, Šamaj J, Chua NH, Barlow PW, Volkmann D. 2000. Root hair formation: F-actin-dependent tip growth is initiated by local assembly of profilin-supported F-actin meshworks accumulated within expansin-enriched bulges. Developmental Biology 227, 618–632.

Baxter A, Mittler R, Suzuki N. 2014. ROS as key players in plant stress signalling. Journal of Experimental Botany 65, 1229–1240.

Beemster GTS, De Vusser K, De Tavernier E, De Bock K, Inze D. 2002. Variation in growth rate between Arabidopsis ecotypes is correlated with cell division and A-type cyclin-dependent kinase activity1. Plant physiology 129, 854–864.

Bus JS, Aust SD, Gibson JE. 1974. Superoxide-and singlet oxygen-catalysed lipid peroxidation as a possible mechanism for paraquat (methyl viologen) toxicity. Biochemical and Biophysical Research Communications 58, 749–755.

Clough SJ, Bent AF. 1998. Floral dip: a simplified method for *Agrobacterium*-mediated transformation of *Arabidopsis thaliana*. The Plant Journal 16, 735–743.

Dat J, Vandenabeele S, Vranová E, Van Montagu M, Inzé D, Breusegem F. 2000. Dual action of the active oxygen species during plant stress responses. Cellular and Molecular Life Science CMLS 57, 779–795.

Daudi A, Cheng Z, O’Brien JA, Mammarella N, Khan S, Ausubel FM, Bolwell GP. 2012. The apoplastic oxidative burst peroxidase in Arabidopsis is a major component of pattern-triggered immunity. Plant Cell 24, 275–287.

Desikan R, Reynolds A, Hancock JT, Neill SJ. 1998. Harpin and hydrogen peroxide both initiate programmed cell death but have differential effects on defence gene expression in Arabidopsis suspension cultures. Biochemical Journal 330, 115–120.

Diet A, Brunner S, Ringli C. 2004. The *enl* mutants enhance the *lrx1* root hair mutant phenotype of *Arabidopsis thaliana*. Plant and Cell Physiology 45, 734–741.

Dietz K-J, Jacob S, Oelze M-L, Laxa M, Tognetti V, Nunes de Miranda SM, Baier M, Finkemeier I. 2006. The function of peroxiredoxins in plant organelle redox metabolism. Journal of Experimental Botany 57, 1697–1709.

Douglas CJ. 1996. Phenylpropanoid metabolism and lignin biosynthesis: from weeds to trees. Trends in Plant Science 1, 171–178.

Dvořák P, Krasylenko Y, Ovečka M, Basheer J, Zapletalová V, Šamaj J, Takáč T. 2020. *In vivo* light-sheet microscopy resolves localisation patterns of FSD1, a superoxide dismutase with function in root development and osmoprotection. Plant, Cell & Environment in press, doi:10.1111/pce.13894

Farrington JA, Ebert M, Land EJ, Fletcher K. 1973. Bipyridylium quaternary salts and related compounds. V. Pulse radiolysis studies of the reaction of paraquat radical with oxygen. Implications for the mode of action of bipyridyl herbicides. Biochimica et Biophysica Acta (BBA) - Bioenergetics 314, 372–381.

Foyer CH, Noctor G. 2013. Redox signaling in plants. Antioxidants & Redox Signaling 18, 2087–2090.

Gilliland LU, Kandasamy MK, Pawloski LC, Meagher RB. 2002. Both vegetative and reproductive actin isovariants complement the stunted root hair phenotype of the Arabidopsis *act2-1* mutation. Plant Physiology 130, 2199–2209.

Han HJ, Peng RH, Zhu B, Fu XY, Zhao W, Shi B, Yao QH. 2014. Gene expression profiles of Arabidopsis under the stress of methyl viologen: a microarray analysis. Molecular Biology Reports 41, 7089–7102.

Haslekås C, Viken MK, Grini PE, Nygaard V, Nordgard SH, Meza TJ, Aalen RB. 2003. Seed 1-Cysteine peroxiredoxin antioxidants are not involved in dormancy, but contribute to inhibition of germination during stress. Plant Physiology 133, 1148–1157.

Hawkes TR. 2014. Mechanisms of resistance to paraquat in plants. Pest Management Science 70, 1316–1323.

Kabsch W, Mannherz HG, Suck D, Pai EF, Holmes KC. 1990. Atomic structure of the actin: DNAse-I complex. Nature 347, 37–44.

Konig J, Muthuramalingam M, Dietz K-J. 2012. Mechanisms and dynamics in the thiol/ disulfide redox regulatory network: transmitters, sensors and tar-gets. Current Opinion in Plant Biology 15, 261–268.

Krieger-Liszkay A, Kós PB, Hideg E. 2011. Superoxide anion radicals generated by methylviologen in photosystem I damage photosystem II. Physiologia Plantarum 142, 17–25.

Kunert KJ, Dodge AD. 1989. Herbicide-induced radical damage and antioxidative systems. In: Boger P, Sandmann G, eds. Target Sites of Herbicide Action, 1st ed., CRC Press, Florida, USA, 49–63.

Lamkemeyer P, Laxa M, Collin V, et al. 2006. Peroxiredoxin Q of *Arabidopsis thaliana* is attached to the thylakoids and functions in context of photosynthesis. The Plant Journal 45, 968–981.

Larkindale J, Knight MR. 2002. Protection against heat stress-induced oxidative damage in Arabidopsis involves calcium, abscisic acid, ethylene, and salicylic acid. Plant Physiology 128, 682–695.

Levine A, Tenhaken R, Dixon R, Lamb C. 1994. H_2_O_2_ from the oxidative burst orchestrates the plant hypersensitive disease resistance response. Cell 79, 583–593.

Liu SG, Zhu DZ, Chen GH, Gao X-Q, Zhang XS. 2012. Disrupted actin dynamics trigger an increment in the reactive oxygen species levels in the *Arabidopsis* root under salt stress. Plant Cell Reports 31, 1219–1226.

Mauch-Mani B, Slusarenko AJ. 1996. Production of salicylic acid precursors is a major function of phenylalanine ammonia-lyase in the resistance of Arabidopsis to *Peronospora parasitica*. Plant Cell 8, 203–212.

Maurino VG, Flügge U-I. 2008. Experimental systems to assess the effects of reactive oxygen species in plant tissues. Plant Signaling & Behavior 3, 923–928.

McDowell JM, Huang SR, McKinney EC, An YQ, Meagher RB. 1996. Structure and evolution of the actin gene family in *Arabidopsis thaliana*. Genetics 142, 587–602.

Meagher RB, McKinney EC, Vitale AV. 1999. The evolution of new structures: clues from plant cytoskeletal genes. Trends in Genetics 15, 278–284.

Mhamdi A, Van Breusegem F. 2018. Reactive oxygen species in plant development. Development 145, dev164376.

Mignolet-Spruyt L, Xu E, Idänheimo N, Hoeberichts FA, Mühlenbock P, Brosché M, Van Breusegem F, Kangasjärvi J. 2016. Spreading the news: subcellular and organellar reactive oxygen species production and signalling. Journal of Experimental Botany 67, 3831–3844.

Miller EW, Dickinson BC, Chang CJ. 2010. Aquaporin-3 mediates hydrogen peroxide uptake to regulate downstream intracellular signaling. Proceedings of the National Academy of Sciences of the United States of Amerika 107, 15681–15686.

Mittler R, Vanderauwera S, Suzuki N, Miller G, Tognetti VB, Vandepoele K, Gollery M, Shulaev V, Van Breusegem F. 2011. ROS signaling: the new wave? Trends in Plant Science 16, 1360–1385.

Mittler R. 2017. ROS are good. Trends in Plant Science 22, 11–19.

Mouratou B, Biou V, Joubert A, Cohen J, Shields DJ, Geldner N, Jürgens G, Melançon P, Cherfils J. 2005. The domain architecture of large guanine nucleotide exchange factors for the small GTP-binding protein Arf. BMC Genomics 6, 20.

O’Brien IEW, Baguley BC, Murray BG, Morris BAM, Ferguson IB. 1998. Early stages of the apoptotic pathway in plant cells are reversible. The Plant Journal 13, 803–814.

Ovečka M, Lang I, Baluška F, Ismail A, Illeš P, Lichtscheidl IK. 2005. Endocytosis and vesicle trafficking during tip growth of root hairs. Protoplasma 226, 39–54.

Pallotta MA, Graham RD, Langridge P, Sparrow DHB, Barker SJ. 2000. RFLP mapping of manganese efficiency in barley. Theoretical and Apllied Genetics 101, 1100–1108.

Pandya-Kumar N, Shema R, Kumar M, et al. 2014. Strigolactone analog GR24 triggers changes in PIN2 polarity, vesicle trafficking and actin filament architecture. New Phytologist 202, 1184–1196.

Park H-J, Miura Y, Kawakita K, Yoshioka H, Doke N. 1998. Physiological mechanisms of a sub-systemic oxidative burst triggered by elicitor-induced local oxidative burst in potato tuber slices. Plant & Cell Physiology 39, 1218–1225.

Pickett CB, Lu AYH. 1989. Glutathione S-transferases: gene structure, regulation, and biological function. Annual Review of Biochemistry 58, 743–764.

Ramel F, Sulmon C, Bogard M, Couée I, Gouesbet G. 2009. Differential patterns of reactive oxygen species and antioxidative mechanisms during atrazine injury and sucrose-induced tolerance in *Arabidopsis thaliana* plantlets. BMC Plant Biology 9, 28.

Rao MV, Davis KR. 1999. Ozone-induced cell death occurs via two distinct mechanisms in *Arabidopsis*: the role of salicylic acid. The Plant Journal 17, 603–614.

Riley D, Wilkinson W, Tucker BV. 1976. Biological unavailability of bound paraquat residues in soil. In: Kaufamn D, Still GG, Paulson GD, Bandal SK, eds. Bound and Conjugated Pesticide Residues, American Chemical Society USA 29, 301–353.

Ringli C, Baumberger N, Diet A, Frey B, Keller B. 2002. ACTIN2 is essential for bulge site selection and tip growth during root hair development of Arabidopsis. Plant Physiology 129, 1464–1472.

Ringli C, Baumberger N, Keller B. 2005. The *Arabidopsis* root hair mutants *der2–der9* are affected at different stages of root hair development. Plant and Cell Physiology 46, 1046–1053.

Staiger CJ, Blanchoin L. 2006. Actin dynamics: old friends with new stories. Current Opinion in Plant Biology 9, 554–562.

Sunkar R, Kapoor A, Zhu J-K. 2006. Posttranscriptional induction of two Cu/Zn superoxide dismutase gene in *Arabidopsis* is mediated by downregulation of miR398 and important for oxidative stress tolerance. The Plant Cell 18, 2051–2065.

Takáč T, Obert B, Rolčík J, Šamaj J. 2016. Improvement of adventitious root formation in flax using hydrogen peroxide. New Biotechnology 33, 728–734.

Takáč T, Šamajová O, Luptovčiak I, Pechan T, Šamaj J. 2017. Feedback microtubule control and microtubule-actin cross-talk in *Arabidopsis* revealed by integrative proteomic and cell biology analysis of *KATANIN 1* mutants. Molecular & Cellular Proteomics 16, 1591–1609.

Takác□ T, S□amajová O, Vadovic□ P, Pechan T, Kos□útová P, Ovec□ka M, Husic□ková A, Komis G, S□amaj J. 2014. Proteomic and biochemical analyses show a functional network of proteins involved in antioxidant defense of the *Arabidopsis anp2anp3* double mutant. Journal of Proteome Research 13, 5347–5361.

Takeda S, Gapper C, Kaya H, Bell E, Kuchitsu K, Dolan L. 2008. Local positive feedback regulation determines cell shape in root hair cells. Science 319, 1241–1244.

Vaahtera L, Brosché M, Wrzaczek M, Kangasjärvi J. 2014. Specificity in ROS signaling and transcript signatures. Antioxidants & Redox Signaling 21, 1422–1441.

Vanderauwera S, Suzuki N, Miller G, et al. 2011. Extranuclear protection of chromosomal DNA from oxidative stress. Proceedings of the National Academy of Sciences of the United States of America 108, 1711–1716.

Vaškebová L, Šamaj J, Ovečka M. 2018. Single-point *ACT2* gene mutation in the *Arabidopsis* root hair mutant *der1-3* affects overall actin organization, root growth and plant development. Annals of Botany 122, 889–901.

Voigt B, Timmers ACJ, Šamaj J, Müller J, Baluška F, Menzel D. 2005. GFP-FABD2 fusion construct allows in vivo visualization of the dynamic actin cytoskeleton in all cells of *Arabidopsis* seedlings. European Journal of Cell Biology 84, 595–608.

Wallström SV, Aidemarka M, Escobar MA, Rasmusson AG. 2012. An alternatively spliced domain of the NDC1 NAD(P)H dehydrogenase gene strongly influences the expression of the ACTIN2 reference gene in *Arabidopsis thaliana*. Plant Science 183, 190–196.

Wang H, Wang S, Lu Y, Alvarez S, Hicks LM, Ge X, Xia Y. 2012. Proteomic analysis of early-responsive redox-sensitive proteins in Arabidopsis. Journal of Proteome Research 11, 412–424.

Wasteneys GO, Galway ME. 2003. Remodeling the cytoskeleton for growth and form: An overview with some new views. Annual Review of Plant Biology 54, 691–722.

Willekens H, Chamnongpol S, Davey M, et al. 1997. Catalase is a sink for H_2_O_2_ and is indispensable for stress defence in C3 plants. The EMBO Journal 16, 4806–4816.

Xiong Y, Contento AL, Nguyen PQ, Bassham DC. 2007. Degradation of oxidized proteins by autophagy during oxidative stress in *Arabidopsis*. Plant Physiology 143, 291–299.

Yao N, Tada Y, Park P, Nakayashiki H, Tosa Y, Mayama S. 2001. Novel evidence for apoptotic cell response and differential signals in chromatin condensation and DNA cleavage in victorin-treated oats. The Plant Journal 28, 13–26.

Yoshida K, Hara S, Hisabori T. 2015. Thioredoxin selectivity for thiol-based redox regulation of target proteins in chloroplasts. The Journal of Biological Chemistry 290, 14278–14288.

Zhou Y, Yang Z, Guo G, Guo Y. 2010. Microfilament dynamics is required for root growth under alkaline stress in Arabidopsis. Journal of Integrative Plant Biology 52, 952–958.

Zwiewka M, Bielach A, Tamizhselvan P, et al. 2019. Root adaptation to H_2_O_2_-induced oxidative stress by ARF-GEF BEN1-and cytoskeleton-mediated PIN2 trafficking. Plant and Cell Physiology 60, 255–273.

